# Automated cell naming reveals reproducible and variable features of ascidian embryogenesis

**DOI:** 10.1101/2025.10.13.681089

**Authors:** Kilian Biasuz, Haydar Jammoul, Benjamin Gallean, Yanis Asloudj, Julien Laussu, Patrick Lemaire, Grégoire Malandain

## Abstract

Ascidians develop with highly reproducible cell lineages, making them ideal models for quantitative comparisons of morphogenesis between individuals and species. Yet, identifying corresponding cells across embryos has so far relied on slow, manual annotation following the Conklin nomenclature, which limits scalability and consistency.

We present an automated framework that assigns cell identities in three-dimensional time-lapse reconstructions of ascidian embryos by transferring names from a reference set of manually curated embryos. The process operates in two steps: an initiation phase, which globally aligns early embryos to establish initial correspondences, and a propagation phase, which propagates names through time and cell divisions by comparing the pattern of contacts each cell forms with its neighbors.

Applied to eight wild-type *Phallusia mammillata* and one *Ascidiella aspersa* embryo, the pipeline assigns consistent names up to stages containing about 700 cells, corrects inconsistencies in earlier datasets, and extends Conklin’s rules to internal tissues beyond gastrulation. Using this unified reference, we quantify natural variability in division timing and orientation, confirming the global robustness and revealing local variability of ascidian morphogenesis. The same framework also demonstrates its use to quantitatively phenotype experimentally perturbed embryos, such as those with inhibited ERK signaling.

This work provides both a validated collection of coherently named ascidian embryos and open-source tools for automated cell identification and phenotypic comparison, establishing a foundation for systematic, quantitative, and evolutionary analyses of animal development.

## 1 Introduction

How a single fertilized egg reproducibly generates a complex multicellular organism remains a central question in developmental biology (Wolpert, 1994). While the underlying gene networks and biophysical processes are increasingly well characterized, understanding how they together ensure reproducible morphogenesis—and how this reproducibility coexists with enough variability to allow adaptation and evolution— requires quantitative comparisons of development across multiple individuals, species, or experimental conditions (Biasuz et al, 2018; Oates et al, 2009).

Such comparisons rely on the ability to identify corresponding cells between embryos, so that equivalent cellular trajectories can be tracked, compared, and statistically analysed (Castro-González et al, 2012; Knowles and Biggin, 2013). Because embryonic cell trajectories exhibit significant inter-individual differences, achieving this correspondence systematically and at scale remains a major challenge in the construction of digital atlases of animal development (Chaudhary et al, 2021).

Some animal embryos, including those of ascidians or nematodes, develop with nearly deterministic lineages and geometries, enabling corresponding cells to be identified across embryos with single-cell precision (Conklin, 1905; Sulston and Horvitz, 1977). This exceptional reproducibility provides a rigorous framework for quantitatively investigating the origins, structure, and consequences of inter-individual variability in an otherwise highly stereotyped developmental system.

Ascidians, simple chordates closely related to vertebrates, are particularly adapted to this challenge (Boutet and Schierwater, 2021). Their embryos are optically transparent, develop with a nearly invariant cell lineage, and fewer than a thousand cells at the onset of tailbud formation (Lemaire, 2009). These features have made them an exceptional model to investigate how genetic and geometric information jointly control cell behaviour during morphogenesis (Di Gregorio and Hadjantonakis, 2006; Veeman and Reeves, 2015). Recent light-sheet imaging and segmentation pipelines have made it possible to reconstruct every cell in a developing ascidian embryo at high temporal resolution, revealing a remarkable conservation of cell geometry and neighbourhood relationships among individuals (Guignard et al, 2020).

The identification of corresponding cells across embryos, a necessary step to integrate multiple reconstructions into a coherent reference atlas, has, however, remained largely manual. Since Conklin’s pioneering nomenclature of ascidian cells in 1905 (Conklin, 1905), each cell has been named according to its lineage and position relative to its sibling, using a syntax that reflects successive cleavage events. Although this convention provides a consistent logical structure, its application beyond early embryogenesis is time-consuming, error-prone, with *ad hoc* decisions that may not always be consistent between embryos. Consequently, most existing datasets remain limited to a handful of manually annotated embryos, restricting the statistical exploration of developmental variability.

Here, we present an automated framework that scales the assignment of cell identities in three-dimensional time-lapse reconstructions of ascidian embryos by transferring names from a reference population of manually curated embryos. It proceeds in two successive steps. In its initiation phase, embryos at equivalent developmental stages are globally aligned to establish the first correspondences between cells. In the subsequent propagation phase, cell names are propagated through time and across cell divisions by comparing the pattern of physical contacts each cell forms with its neighbors. This combination of global alignment and local relational comparison ensures that naming remains robust to small geometric differences and to natural variability in cell division timing or orientation.

We first applied this strategy to a collection of seven published wild-type *Phallusia mammillata* embryos (Guignard et al, 2020). We systematically identified and corrected inconsistencies present in this earlier datasets. We further extended the Conklin rules to internal tissues beyond gastrulation, allowing the complete naming of mesodermal and endodermal lineages. The ensuing coherent reference atlas names every cell up to initial tailbud stages, containing about 700 cells. This atlas identifies several issues in the published literature and solves longstanding ambiguities in the description of ascidian embryonic lineages.

We then used this atlas to name *de novo* a novel *Phallusia mammillata* embryo and show that the procedure is sufficiently robust to name an embryo from an evolutionary distant ascidian species, *Ascidiella aspersa*, demonstrating the evolutionary generality of the framework.

Finally, this unified reference made it possible, for the first time, to quantitatively assess natural developmental variability across multiple ascidian embryos. Despite the well-known stereotypy of ascidian development, we detect reproducible yet measurable differences in division timing, orientation, and neighborhood topology—confirming the global robustness of the developmental program while revealing its local variability. The same approach can be applied to embryos subjected to experimental perturbations, such as inhibition of ERK signaling.

Together, these developments establish a scalable and reproducible strategy for integrating, comparing, and analyzing digital reconstructions of ascidian embryos. Beyond providing a resource for the ascidian community, this work illustrates how automated cell identity transfer can transform the study of morphogenesis—from descriptive reconstructions of individual embryos to quantitative, comparative analyses of development across individuals, species, and experimental conditions.

## 2 Results

### 2.1 The Conklin cell nomenclature and its limitations

Ascidian embryonic cells are traditionally named according to the nomenclature introduced by Conklin in 1905 (Conklin, 1905) and still used throughout the community. This system combines a hierarchical syntax, which defines how cell names are structured, with a set of decision rules that determine how names are assigned to daughter cells after each division.

Each cell is assigned a unique identifier of the form Fr.p (for example, A6.1), where F denotes the founding lineage at the 8-cell stage (A, a, B, or b for the anterior/posterior and vegetal/animal quadrants), r is the rank of the mitotic cell cycle (e. g. r=3 means that the cell is in its 3rd mitotic cell cycle since fertilization), and p is an integer reflecting the position of the cell relative to its sibling. Because ascidian embryos are bilaterally symmetrical, corresponding cells on the right and left sides share the same name, distinguished by a suffix: right-side cells (from a vegetal view) receive a “*” and left-side cells a “_” (for example, A6.1 and A6.1*) (Figure 1). Conklin’s syntax then specifies that cell Fr.p divides into two daughters named F(r+1).(2p) and F(r+1).(2p–1).

**Fig. 1:**
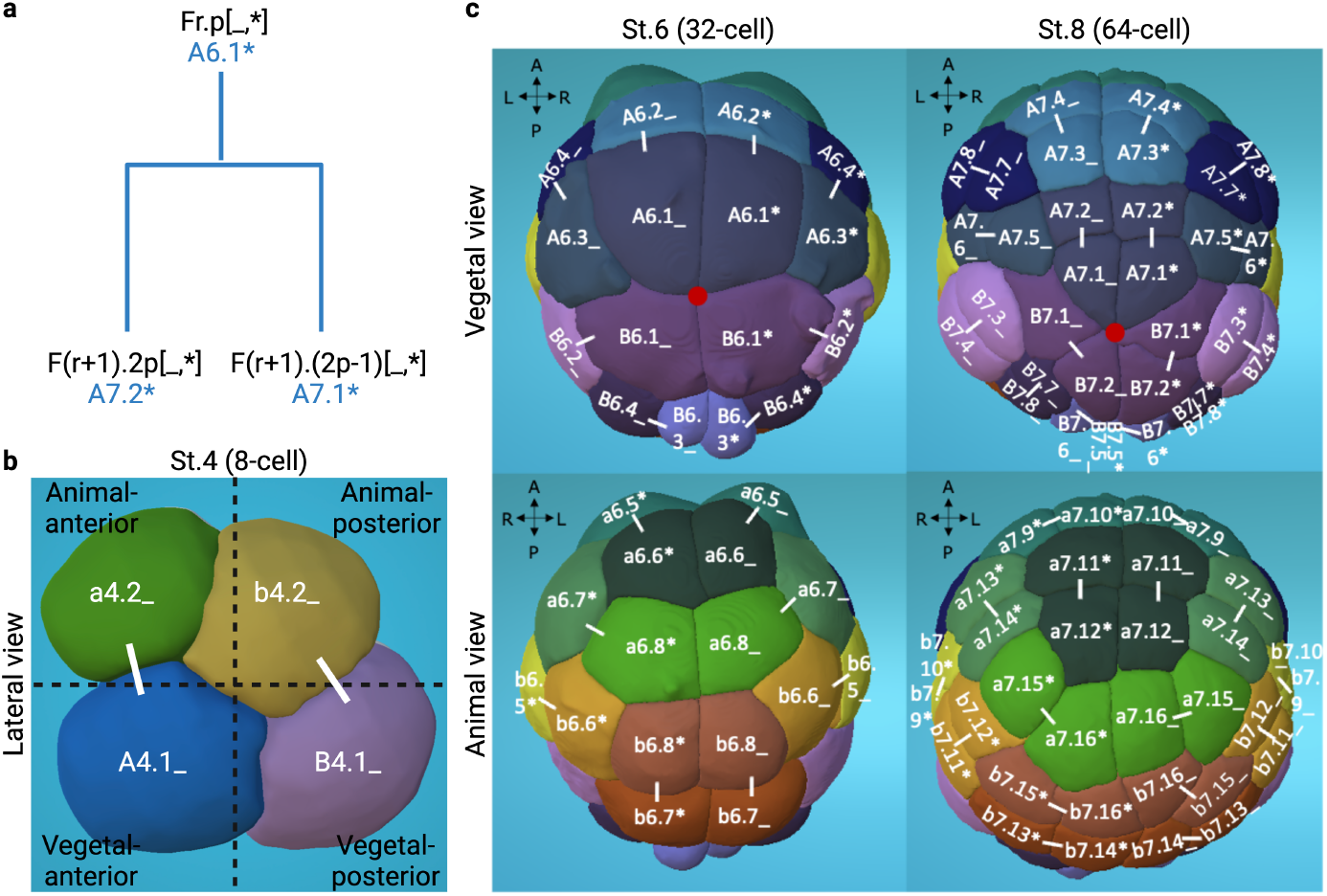
Cell naming of early ascidian embryos according to the Conklin nomenclature. a) Conklin naming scheme, exemplified with the case of the A6.1 cell lineage. b) Lateral view of an 8-cell embryo, showing the position of the 4 founding cell lineages. White lines link sister cells. c) Vegetal (top) and animal (down) views of a 32-cell (left) and a 64-cell (right) stage *Phallusia mammillata* embryo. Each bilateral cell pair at the 32-cell stage is assigned a unique color, which is propagated at the 64-cell stage. Names (in white) are according to Conklin. White lines link sister cells. Antero-Posterior (AP) and Left-Right (LR) axes are represented. The red disks materialize the vegetal pole of the embryo, used as a reference for the measurement of the geodesic distance for the Conklin naming. Snapshots of the Astec-Pm-Stages embryo (https://morphonet.org/dataset/1062) at the indicated stage (Guignard et al, 2020) are visualized using MorphoNet (Gallean et al, 2025; Leggio et al, 2019).

The assignment of one of these these names to each of the daughters follows a geometric decision rule: the daughter whose apical surface lies closer to the vegetal pole of the embryo—measured along the embryonic surface receives the F(r+1).(2p–1) identifier (Conklin, 1905). This rule relies on the notion of geodesic distance to the vegetal pole along the embryo’s external surface and therefore applies only to stages in which all cells remain in contact with the exterior. From the 76-cell stage onward, some divisions occur perpendicular to the embryo surface, producing internal cells whose geodesic distance to the vegetal pole cannot be defined. As a result, internal cells such as the cardiogenic precursors (Stolfi et al, 2010) cannot be named individually within the Conklin system—their conventional designations reflect their larval fates rather than their clonal origins. Likewise after the vegetal pole has fully invaginated, it cannot be used as a reference point and ad-hoc Conklin-inspired rules have to be designed, as was done for the neural plate cells (Cole and Meinertzhagen, 2004; Nicol and Meinertzhagen, 1988a,b).

Thus, while the Conklin naming syntax ensures the consistency of cell names and is broadly applicable beyond the gastrula stages, its decision rules cannot be strictly applied beyond the onset of gastrulation. This makes the Conklin scheme impossible to implement directly in a programmatic way. This limitation motivated us to develop an alternative strategy in which cell identities are transferred from reference embryos rather than calculated from geometric rules.

### 2.2 Cell naming by global registration of cell positions between early embryos

We first tested whether homologous cells between embryos of equivalent developmental stages can be identified on the basis of the conservation of their global spatial positions across embryos. To do so, the embryo to be named was temporally aligned with a reference embryo to identify equivalent developmental stages (see Supp. Methods B.5.1). Embryo sizes were then normalized, and cell barycenters were spatially aligned by affine transformation (Michelin et al, 2015a). Finally, names were transferred by assuming that close barycenters correspond to homologous cells (Figure 2a; see Supp. Methods B.6.1).

**Fig. 2:**
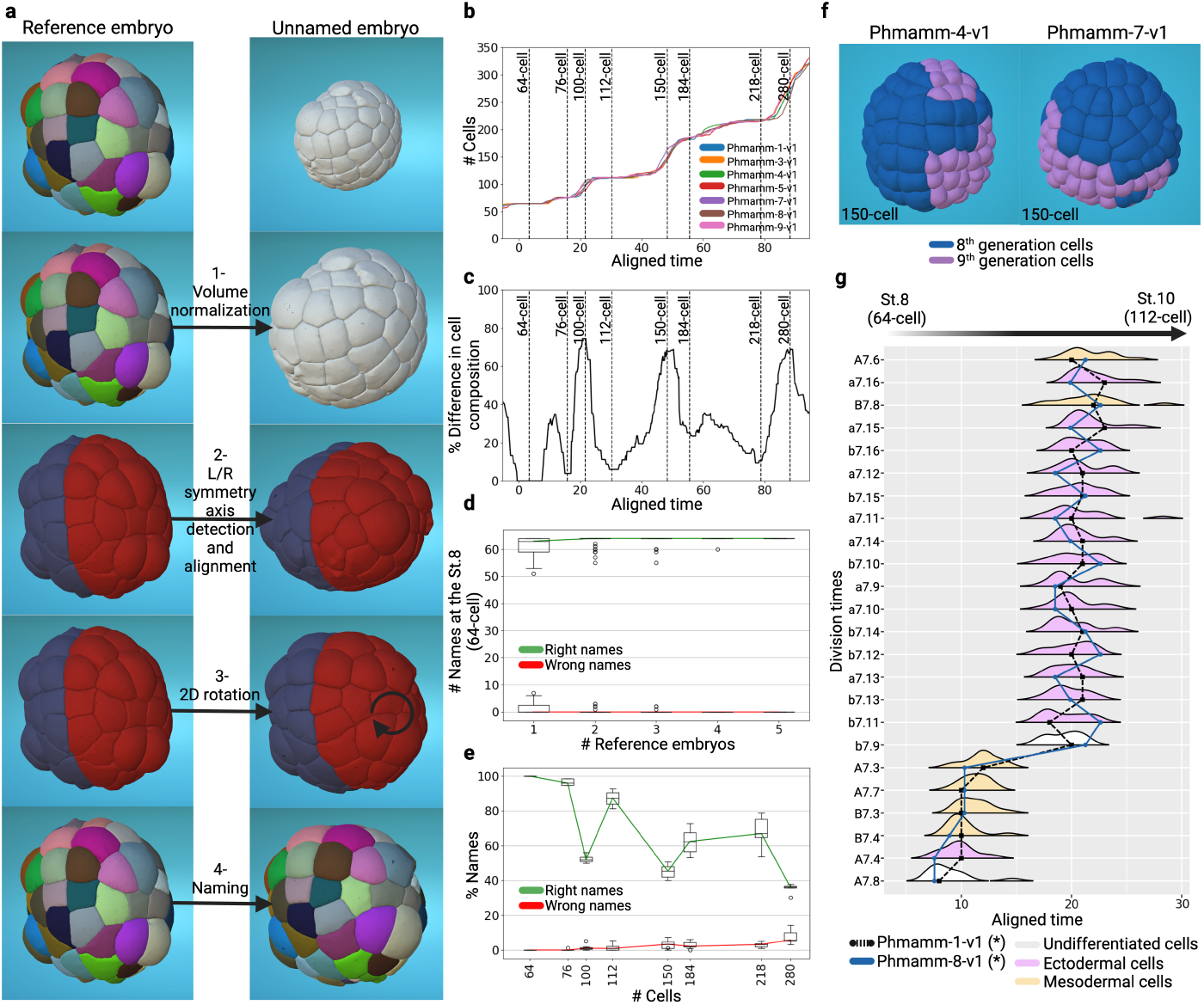
Assessment and determinants of the efficiency of cell name transfer by co-registration with named embryos. a) The 4 steps of the identification process of homologous cells between two embryos. b) Plot of the number of cells as a function of developmental time for the 7 reference embryos. c) Differences in cell composition across the reference embryos. Purple horizontal lines indicate the position of waves of epidermal cell division in Phmamm-1-v1. d) Number of correctly and erroneously named cells at the 64-cell stage with respect to the number of used reference atlases. Top, middle, and bottom lines in the box plots indicate 75^*th*^ percentile, median, and 25^*th*^ percentile of data, respectively. Each reference embryo is tested with the exception of Phmamm-9-v1, which starts with 72 cells. e) Fraction of correctly and erroneously named cells at different time points (identified by cell counts): each reference embryo is tested against the other ones (Phmamm-9-v1 being discarded for the 64-cell stage). f) Visualization using MorphoNet of differing cell division orders in the epidermis of the Phmamm-4-v1 and Phmamm-7-v1 embryos at 150-cells. 8^*th*^-generation cells are in blue, 9^*th*^-generation cells in purple. g) Representation of the distributions of the division times of indicated cells across the embryo collection during the transition from the 64-to 112-cell stages). The dotted black and blue lines correspond to the right instances of the indicated cell divisions in embryos Phmamm-1-v1 and Phmamm-8-v1. Note that the lines cross frequently.

We evaluated this procedure by renaming each previously published (Guignard et al, 2020) reference embryo (Phmamm-1/3/4/5/7/8-v1) at the 64-cell stage through transfer from the others. Over these 30 experiments, an average of 61 cells per embryo (median: 63 cells) were correctly named, with up to 6 unnamed and 7 misnamed cells in the least accurate cases (see Figure 2d).

These residual errors most likely arose from geometric variability that limits the accuracy of affine co-registration of pairs of embryos. Accordingly, averaging over multiple named embryos (see Supp. Methods B.6.2) effectively reduced the numbers of unnamed and misnamed cells (Figure 2d). Using five references sufficed to correctly rename all cells of each Phmamm reference embryo at the 64-cell stage.

At later developmental stages, we expected naming accuracy to decline as cell numbers increased. Instead, naming efficiency varied non-monotonically across stages, with lower accuracy at the 100-, 150-, and 280-cell stages and higher accuracy at the 76-, 112-, and 218-cell stages (Figure 2e). This unexpected jigsaw pattern closely mirrored changes in cellular composition between embryos, which were greater than expected for embryos traditionally described as having invariant cell lineages: at the 100-, 150-, and 280-cell stages, corresponding to rounds of epidermal tissue divisions, fewer than 25% of cells were shared across all six embryos (Figure 2c). These parallel stage-dependent fluctuations in cell composition and naming accuracy arose from changes in the order of cell divisions within each dividing tissue, caused by small cell division heterochronies (Figure 2f,g).

In summary, cell naming by transfer from a population of temporally and spatially co-registered embryos provides an efficient strategy to systematically name embryonic cells *de novo* at the 64-, 76-, and 112-cell stages. Slight heterochronies across embryos, in particular during epidermal cell divisions, however, limit the usefulness of a global registration strategy for naming cells at later developmental stages.

### 2.3 Local cell–cell contacts discriminate sibling cells in

#### *Phallusia* embryos

To extend automated naming beyond this initial stage, we introduced a propagation step, in which names are transferred across successive divisions within each embryo. Given that local cell–cell contact patterns are expected to be less sensitive to geometric variability than global positions, we based the propagation step primarily on this local information (Figure S-11b).

For local contact information to be useful in naming cells, neighborhood relationships (see glossary A) must be informative, stable over a cell’s lifetime, conserved between embryos, and distinct between sister cells.

Figure 3a shows that between the cleavage and neurula stages in the 7 reference Phmamm embryos (Guignard et al, 2020), each embryonic cell established significant physical contacts (*>* 5% of its surface) with an average of 5 to 6 immediate neighbors.

**Fig. 3:**
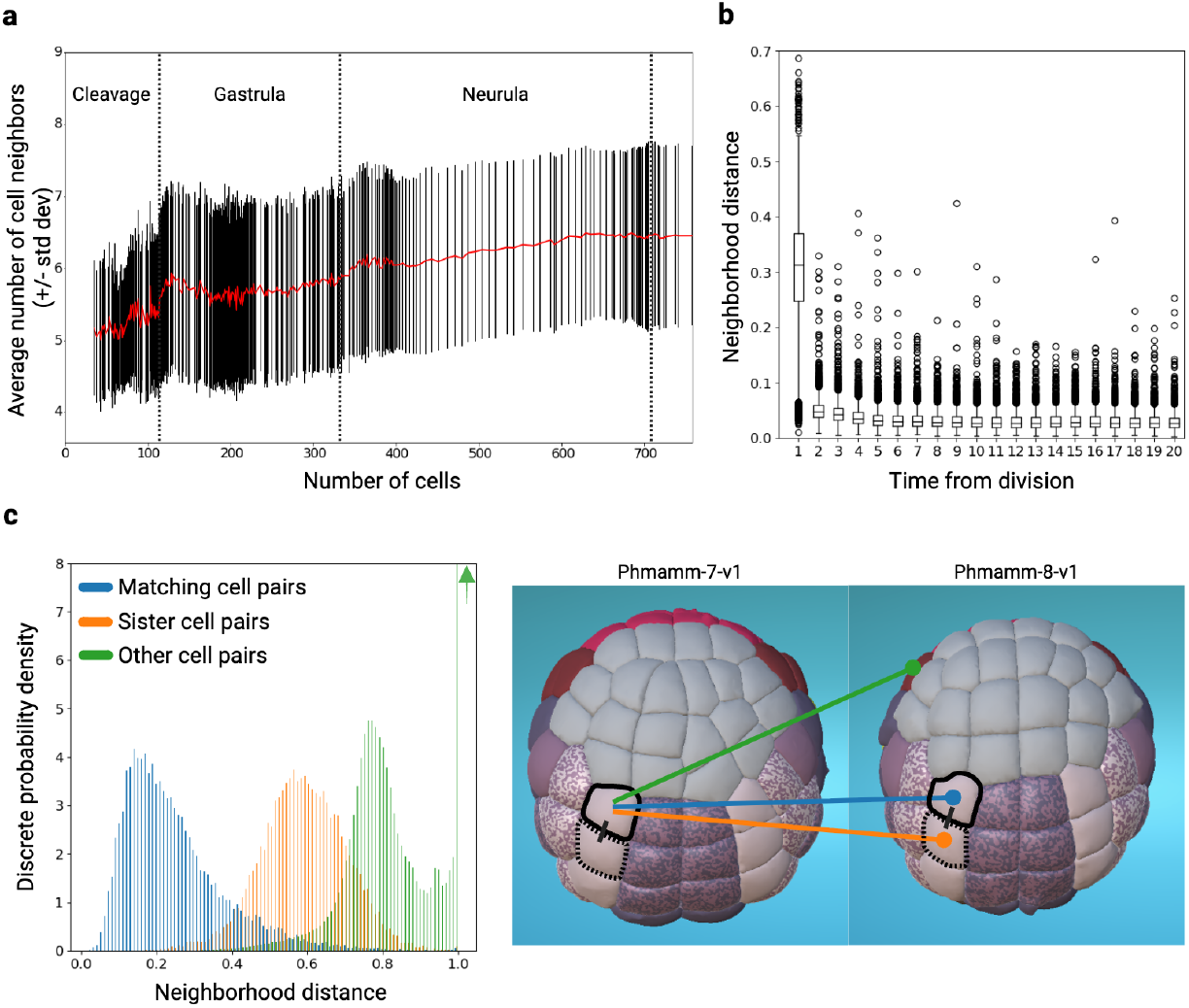
Temporal and spatial reproducibility of cell neighborhoods during *Phallusia mammillata* development. a) Evolution of the average number (red line) of cell neighbors touching a cell over *>*5% of its surface (+/-standard deviation, black) during early *Phallusia mammillata* development. b) Stability of neighborhood analysis. For each cell, the stability of its neighborhood was determined by calculating the neighborhood distance between two successive time points throughout the entire lifetime of the cell. Top, middle, and bottom lines in the box plot indicate 75^*th*^ percentile, median, and 25^*th*^ percentile of data, respectively. c) Left: neighbor-hood distance measurements between pairs of cells: matching cells (blue), sister cells (orange), other cells (green, includes all cells of the same generation except matching and sister cells) within and across embryos. The green arrow indicates that within the limits of the graph shown, only part of the height of the discrete probability density for the neighborhood distance class with a value of 1.0 is visible (total height=12.7). Right: schematic illustrating for a given cell in Phmamm-7-v1 (outlined in black) the different types of cell pairs from which the neighborhood distances are calculated. The cell surrounded by black dotted lines corresponds to the sister cell in this example. Vegetal views of Phmamm-7-v1 and Phmamm-8-v1 at the 112-cell stage (St. 10, initial gastrula). Cells are color-coded according to their larval fate (see Figure S-14). a-c) n=7 embryos.

When comparing local neighborhoods across embryos, slight asynchronies in cell-division timing introduce inconsistencies: while in embryo *E* a cell *i* may contact its neighbor *j*, in another embryo *E*′, *j* has already divided and *i* contacts its two daughters. To compensate for these asynchronies, we defined a common virtual neighborhood in which cells from equivalent lineage generations can be compared directly. In this common framework, the contact area between cells *i* and *j* in *E*′ is computed as the sum of *i*’s contacts with the daughters of *j* (Figure S-12a).

In this unified reference frame, the neighborhood of cell *i* in embryo *E* at time *t*, noted *c*_*E*_ (*i*; *t*), is represented by the vector of its normalized contact areas with all directly adjacent cells at this time point. The distance between the neighborhoods between two cells is then defined as the fraction of the total contact areas made with different neighbors (see Supp. Methods B.7.1). This measure can be computed for any pair of cells, whether they belong to the same or to different embryos, and at identical or distinct time points.

Using this neighborhood distance, we first analyzed the temporal stability of each cell’s local neighborhood across successive time points (Figure 3b, Figure S-13). Neighborhood distances increased sharply immediately after each cell division but stabilized soon thereafter (Figure 3b), consistent with our previous analysis (Guignard et al, 2020). In all subsequent analyses, contact areas computed 4 time points after division were considered representative of the stable contacts maintained by a cell during most of its lifetime.

We next assessed neighborhood distances across different classes of cell pairs. As shown in Figure 3c, the distributions of distances between matching cells, sister cells, and all other pairs of cells of the same generation—within and across embryos—showed minimal overlap. As expected, neighborhoods of unrelated cells were the most dissimilar. On average, matching cells contacted different neighbors on about 20% of their surface area, consistent with the high reproducibility of cell contacts reported during ascidian embryogenesis (Guignard et al, 2020). By contrast, sister cells differed over roughly 60% of their contact surface.

Taken together, these results demonstrate that contact areas between cells stabilize shortly after division, are highly conserved between embryos, and differ markedly between sister cells. The prerequisites for using local neighborhood information to identify individual cells and distinguish them from their sisters are therefore satisfied.

### 2.4 Propagation of cell names through local contact information

We next explored whether this information can be used to propagate cell names across cell divisions (Figure 4). We start from a situation where a cell Fr.p of a new embryo *E* divides to produce two daughters that we wish to name by transfer from a reference embryo *R* (Figure 4). We assume that Fr.p has also divided in embryo *R* to produce two daughters named F(r+1).(2p) and F(r+1).(2p-1), following the Conklin syntax.

**Fig. 4:**
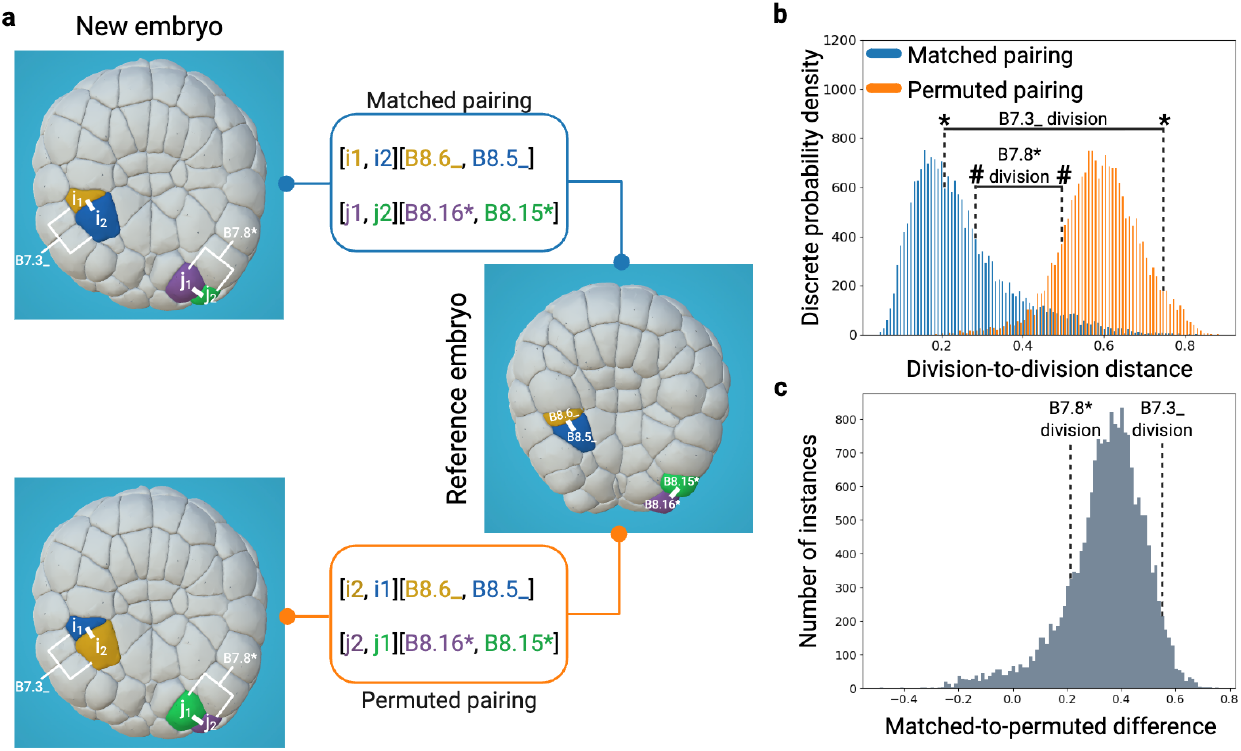
Distinguishing matched and permuted sister cell pair names. a) Illustrations of the matched (top) and permuted (bottom) pairing of the daughters of B7.3 or B7.8* between a test unnamed embryo (Phmamm-1-v1) and a named reference embryo (Phmamm-8-v1). Colors indicate the ordering of homologous sister cell pairs used to compute division distances. b) Histograms of the pairwise division distances between homologous matched (blue) or permuted (orange) sister cell pairs across the 7 reference embryos. (*) and (#) highlight the positions on the histograms of the matched or permuted division distance values for B7.3 (*) and B7.8* (#) divisions between Phmamm-1-v1 and Phmamm-8-v1. c) Histogram of the computed differences between permuted and matched division distances. The positions on the histogram of the division distance differences values for the B7.8* division (value=0.22) and the B7.3 division (value=0.54) are indicated.

Naming the two daughter cells *c*_*E*_ (*i*_1_) and *c*_*E*_ (*i*_2_) of Fr.p in embryo *E* by reference to *R* is therefore equivalent to determining which of the two possible ordered cell pairs, [*c*_*E*_ (*i*_1_), *c*_*E*_ (*i*_2_)] and [*c*_*E*_ (*i*_2_), *c*_*E*_ (*i*_1_)], spatially matches the ordered pair [F(r+1).(2p), F(r+1).(2p-1)] in *R*. This matching process is illustrated in Figure 4a,b for the divisions of cells B7.3 and B7.8*.

To identify the correct pairing of daughter cells between embryos E and R, we introduced a division distance (see Supp. Methods B.7.2), extending the neighborhood distance to cell pairs. This metric is expected to yield lower values for correctly matched homologous cell pairs than for permuted pairs, ensuring reliable name transfer (Figure 4).

To test this hypothesis, we analyzed the distributions of division distances between matched and permuted homologous pairs across the population of seven named reference embryos. As expected, matched cell pairs exhibited significantly lower distances (mean: 0.25, standard deviation: 0.12) than permuted pairs (mean: 0.59, standard deviation: 0.10) (Figure 4b). Furthermore, the difference in distances between homologous matched and permuted cell pairs showed a high mean (0.34) and a low standard deviation (0.15) (Figure 4c).

Taken together, these results suggest that homologous daughter cells can be robustly paired across embryos, enabling the propagation of names across a cell division.

### 2.5 End-to-end validation of automated cell naming and harmonisation of reference datasets

We next applied our complete automated naming pipeline, combining global initiation and local propagation, to rename the seven segmented Phmamm embryos *de novo* from the 64-cells stage onward by automated transfer from Phmamm-8-v1, as previously done manually (Guignard et al, 2020). This analysis served two purposes: first, to validate that the automated procedure reliably reproduces manual namings; and second, to use it as a benchmark to identify and correct potential inconsistencies in the reference manually named dataset.

The automated and manual procedures agreed on the names of 86.3-95.9% of cells, depending on the embryo (Table 1), demonstrating that our iterative procedure correctly names a large majority of cells.

**Table 1:**
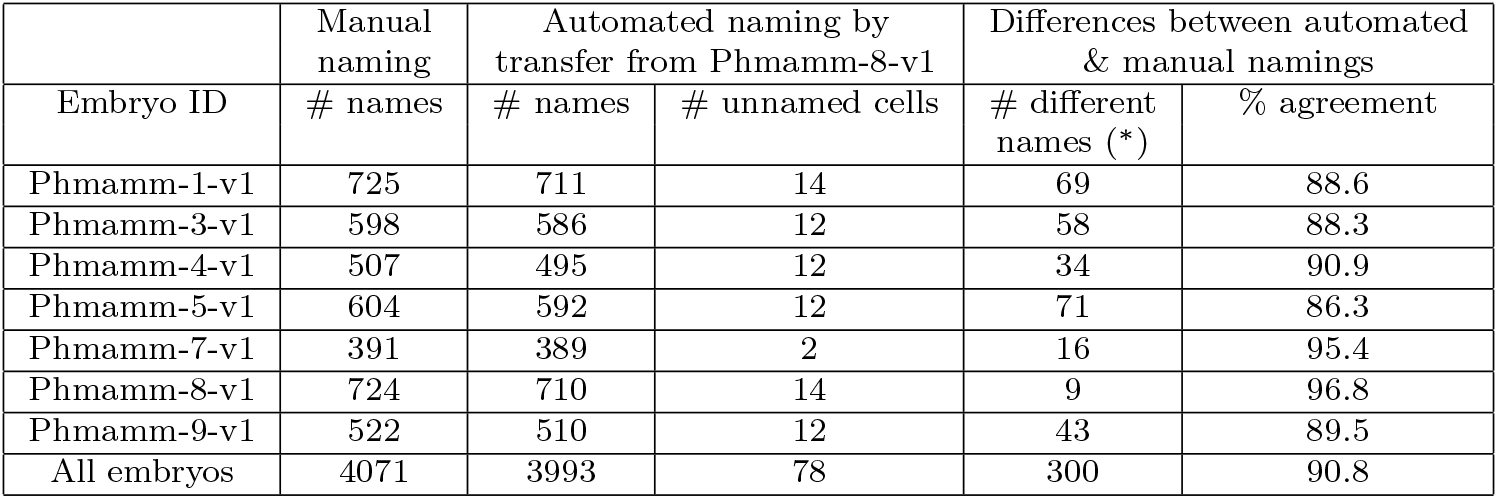
Comparison of the manual and automated *de novo* naming of Phmamm embryos by transfer from Phmamm-8-v1. (#)=“Number of”. (*): Includes both primary and induced differences (see glossary A).

Having validated the overall performance of the automated naming, we next sought to understand and resolve discrepancies between the original manual and automated namings (5–15% of cells). The negative division distance values observed in Figure 4c, raised the possibility that part of the original manual naming was erroneous. We indeed identified three main sources of errors: violation of the Conklin syntax, segmentation or lineage-reconstruction errors, and permutations between sister cell names.

Parsing the lineage and naming information (see Methods 5.4) identified 36 cell names that did not respect the Conklin syntax (Table C1). For example, the left b10.91 cell in Phmamm-1-v1 was mistakenly named as its right counterpart, b10.91*, while the daughters of b9.58* in Phmamm-4 were incorrectly named b10.113* and b10.114*, instead of the expected b10.115* and b10.116*. This analysis also uncovered a few segmentation or lineage-reconstruction errors, which were corrected.

A more frequent problem was the inversion of sister-cell names. Figure 5a,b illustrates one such case, where the daughters of b7.12 cells are permuted between Phmamm-8-v1 and Phmamm-5-v1. To objectively detect such inconsistencies, we compared both the original and the permuted versions of each division across all embryos using the division distance metric (Figure 5c). In this example, all embryos showed consistent division-distance patterns except Phmamm-8-v1, indicating that the names of the b7.12 daughters in Phmamm-8-v1 need to be inverted.

**Fig. 5:**
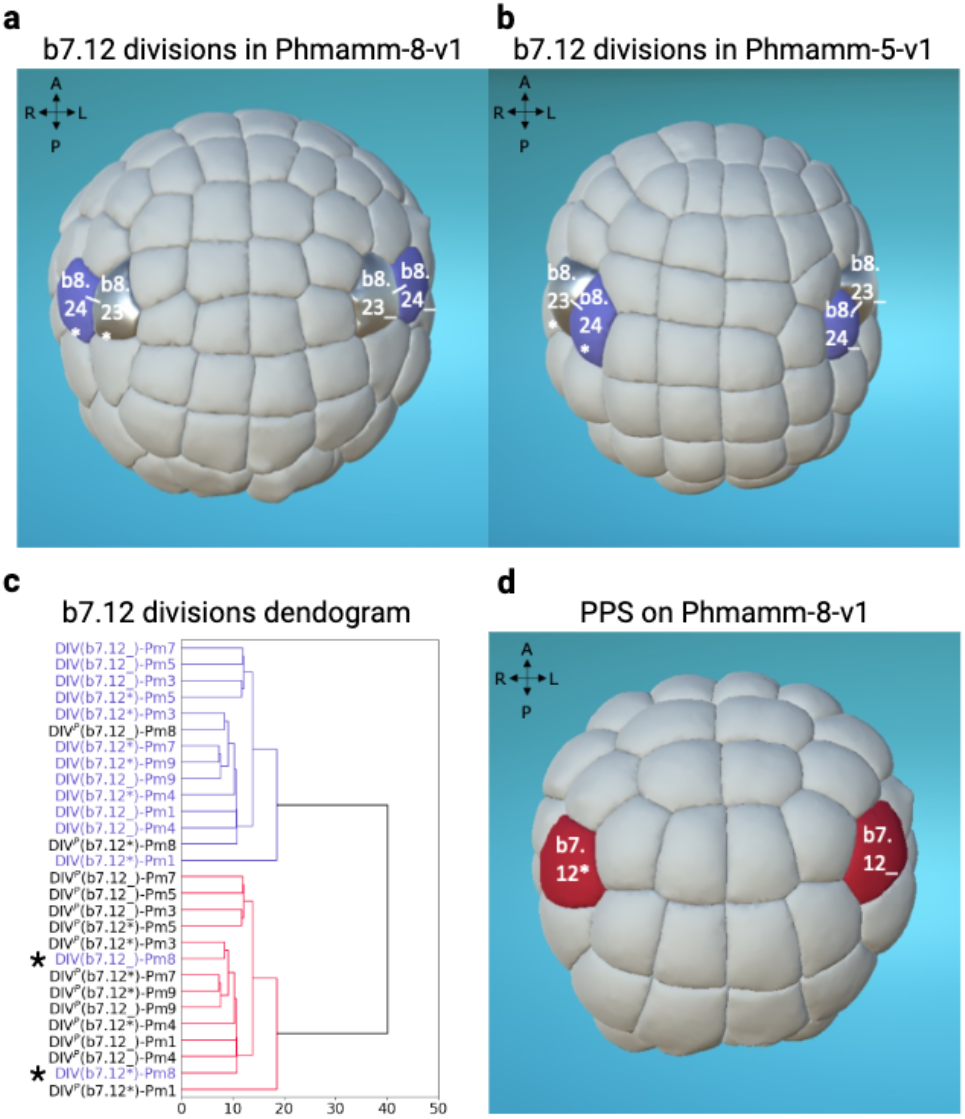
Systematic identification of sister cells with permuted manual names in a minority of embryos. a-b) Visualization using MorphoNet (Leggio et al, 2019) of manually assigned names for daughters of b7.12 and b7.12* divisions in Phmamm-8-v1 (a) and Phmamm-5-v1 (b). Cell names are displayed and cells with the same name between the two embryos are color coded. c) Dendrogram of b7.12 divisions from the 7 manually named Phmamm-v1 embryos (referred to as DIV(b7.12*/_-Pm) or permuted (DIV^*P*^ (b7.12*/_-Pm) with the division distance as a metric. Asterisks (*) highlight the positions of the original manual naming of the b7.12 divisions of Phmamm-8-v1 in the dendrogram. d) Visualization of the b7.12 cells with a positive value of the permutation proposal score (PPS, color-coded in red) on Phmamm-8-v1 using MorphoNet. a-b-d) Animal views, Antero-Posterior (AP) and Left-Right (LR) axes are represented.

Extending this approach to all divisions, we introduced a Permutation Proposal Score (PPS, see Methods 5.5) to quantify the likelihood of a naming inversion. The PPS is the signed difference between (i) the summed division distances of an unpermuted division of the tested embryo and those of all others, (ii) the same sum computed for the permuted version of the division. A positive PPS value therefore indicates that the permuted configuration is more consistent with the other embryos, suggesting that a naming inversion occurred.

Across the dataset, fifty-four divisions showed positive PPS values. After visual analysis in MorphoNet (Figure 5b), all proposed permutations were accepted to maximize the consistency of the namings between embryos. These included clear manual errors, as well as ambiguous cases involving internal cells or atypical division angles, for which adopting the consensus configuration maximised naming consistency. In total, 108 sister-cell names (two per division) were corrected, which in turn propagated to 42 granddaughter names whose mother names were permuted (Table C1).

These corrections notably affected the original Phmamm-8-v1 dataset, which had served as a reference for manual naming of other embryos in (Guignard et al, 2020). In total, 38 cell names in Phmamm-8-v1 were adjusted, leading to the corrected reference Phmamm-8-v2. Reapplying the automated naming procedure to the other six reference embryos using Phmamm-8-v2 as a reference significantly improved the agreement between automated and corrected reference namings (Table 2).

**Table 2:**
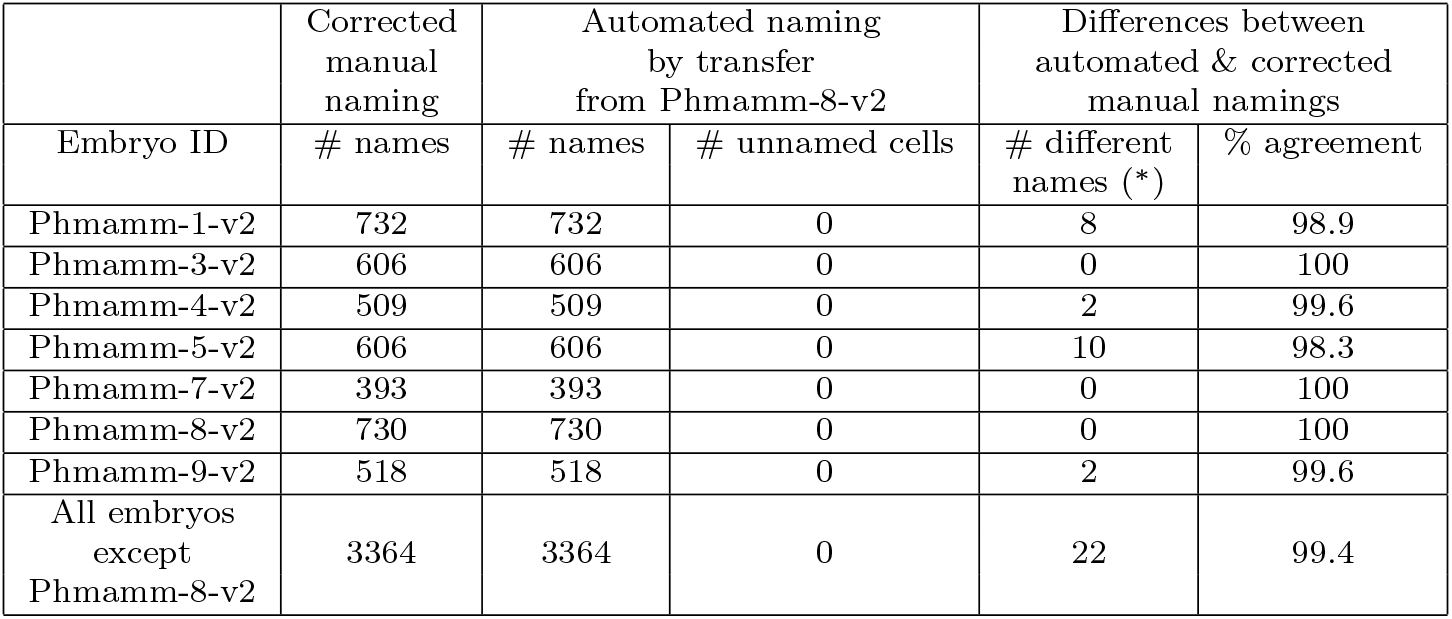
Comparison of the corrected manual and *de novo* automated naming of Phmamm embryos by transfer from the corrected Phmamm-8-v2. (#)=“Number of”. (*): includes both primary and induced differences (see glossary A).

Overall, these corrections maximised the internal consistency and reliability of cell naming across the entire dataset.

### 2.6 Naming with respect to a collection of reference embryos increases accuracy and robustness to inter-individual variation

Residual discrepancies between automated and corrected manual namings point to genuine inter-individual differences in embryonic geometry, notably in cell division orientation, which could make automated naming sensitive to the choice of reference embryo. To evaluate this potential reference dependence, we re-applied the automated naming process to the six remaining Phmamm embryos, this time using Phmamm-5 as the corrected reference (Table 3). As illustrated in Figure S-16, differences in cleavage orientations, for example in the A7.2 cells between Phmamm-8-v2 and Phmamm-5-v2, produced distinct namings of the A7.2 daughters in Phmamm-3, depending on which reference was used. These results confirm that naming by transfer from a single embryo is sensitive to inter-individual geometric variation.

**Table 3:** Comparison of the corrected manual and *de novo* automated naming of Phmamm embryos by transfer from the corrected Phmamm-5-v2. (#)=“Number of”. (^$^)=Cells that are not named because they do not exist and are not named in the reference embryo, here Phmamm-5-v2. These unnamed cells are therefore not counted as differences between automated and manual naming. (*): Includes both primary and induced differences (see glossary A).

|  | Corrected manual naming | Automated naming by transfer from Phmamm-5-v2 |  | Differences between automated & corrected manual namings |  |
| --- | --- | --- | --- | --- | --- |
| Embryo ID | # names | # names | # unnamed cells (§) | # different names (*) | % agreement |
| Phmamm-1-v2 | 732 | 620 | 112 | 22 | 96.5 |
| Phmamm-3-v2 | 606 | 590 | 16 | 16 | 97.3 |
| Phmamm-4-v2 | 509 | 509 | 0 | 4 | 99.2 |
| Phmamm-5-v2 | 606 | 606 | 0 | 0 | 100 |
| Phmamm-7-v2 | 393 | 393 | 0 | 4 | 99 |
| Phmamm-8-v2 | 730 | 620 | 110 | 12 | 98.1 |
| Phmamm-9-v2 | 518 | 516 | 2 | 14 | 97.3 |
| All embryos except Phmamm-5 | 3488 | 3248 | 240 | 72 | 97.9 |

To overcome this limitation, we extended the automated procedure to use as reference a population of embryos *R* rather than a single individual. Cell names in an embryo *E* were assigned based on the *mean division distance* between each division in *E* and its homologs in *R* (see Supp. Methods B.7.3). Automated *de novo* renaming of each Phmamm embryo by transfer from the population of all other reference embryos achieved higher accuracy (Table 4) than using either Phmamm-8-v2 (Table 2) or Phmamm-5-v2 (Table 3) alone. The multi-reference approach reached 99,9% agreement between automated and corrected reference namings, including divisions with variable orientations, such as that of A7.2 (Figure S-16).

**Table 4:** Comparison of the manual corrected and automated *de novo* naming of Phmamm embryos by transfer from the corrected reference population (except the tested one). (#)=“Number of”. (*): Includes both primary and induced differences (see glossary A).

|  | Manual naming | Automated naming by transfer from the population |  | Differences between automated & corrected manual namings |  |
| --- | --- | --- | --- | --- | --- |
| Embryo ID | # names | # names | # unnamed cells | # different names (*) | % agreement |
| Phmamm-1-v2 | 732 | 732 | 0 | 2 | 99.7 |
| Phmamm-3-v2 | 606 | 606 | 0 | 0 | 100 |
| Phmamm-4-v2 | 509 | 509 | 0 | 0 | 100 |
| Phmamm-5-v2 | 606 | 606 | 0 | 2 | 99.7 |
| Phmamm-7-v2 | 393 | 393 | 0 | 0 | 100 |
| Phmamm-8-v2 | 730 | 730 | 0 | 0 | 100 |
| Phmamm-9-v2 | 518 | 518 | 0 | 0 | 100 |
| All embryos | 4094 | 4094 | 0 | 4 | 99.9 |

Together, these results show that using a population of reference embryos markedly enhances both the accuracy and the robustness of automated naming, providing a consistent framework for quantitative comparisons despite inter-individual variability in cleavage geometry.

### 2.7 Expanding Conklin rules beyond the onset of gastrulation

The consistent naming of all cells across embryos revealed that all cells, even internally located and therefore previously excluded from Conklin’s scheme, occupy reproducible positions up to the early tailbud stage. This harmonized dataset therefore provides the foundation for a systematic extension of the Conklin rules to internal cells up to the 10^*th*^ cell generation.

Internal tissues were defined as encompassing all embryonic tissues except the epidermis, which remains external, and the neural plate, for which cell names were already established up to the 10^*th*^ generation (Cole and Meinertzhagen, 2004; Nicol and Meinertzhagen, 1988a,b). From the 8^*th*^ generation onward, several cells are not named in Conklin’s original work (Conklin, 1905), as some divisions produce internal daughters and the vegetal pole used as a geometric reference becomes internalized during gastrulation. We therefore extended Conklin’s system from this generation onward, up to the 10^*th*^, covering a total of 98 cells previously lacking a defined name.

Because the orientation of certain divisions varies between embryos, we first determined for each division the most frequent orientation observed across the seven Phmamm reference embryos. For cells that retained external contact during gastrulation, names were assigned following Conklin’s original rule: when both daughters remained in contact with the exterior, the cell closer to the vegetal pole received the lower index; when only one did, that cell received the lower index. As an example, this logic led to invert the names of the two daughters of A7.6 in the reference collection, to place A8.11 in contact with the exterior. In total 47 cells were named according to this strategy.

When both sister cells were internal, we extended Conklin’s logic by virtually “peeling” away superficial cell layers, using the MorphoNet 3D anatomical browser (see Methods 5.6). In the resulting partial embryos, the relevant internal cells again came into contact with the simulated exterior, allowing us to measure their geodesic distance to the vegetal pole. When this distance provided an unambiguous ordering, the lower index was assigned to the cell closer to the vegetative pole. For divisions where the axis of orientation made this rule ambiguous, we adopted the predominant orientation observed across embryos, giving precedence—by order—to the more medial, anterior, or dorsal sister. This approach allowed to name 49 internal cells, including 8^*th*^ and 9^*th*^ generation cells in the B7.5 cardiopharyngeal lineage, previously designated by fate rather than by Conklin-type naming (Razy-Krajka et al, 2014) (Figure S-15).

Beyond these general rules, one particular case required special attention. The division of b8.17, a descendant of b6.5, produces two external daughters, b9.33 and b9.34, that were initially unnamed by Conklin (Conklin, 1905) and have since been inconsistently described in the literature, with opposite naming conventions not only between ascidian species but even within a single species as observed in *Ciona* (Hudson and Yasuo, 2005; Imai et al, 2006; Noda et al, 2008). Because of this long-standing ambiguity, we revisited this division in detail. According to our extended rules, the cell closer to the vegetal pole should receive the lower index. However, to ensure consistency with the reference nomenclature established in *Halocynthia* (Nishida, 1986), we adopted his convention, naming the smaller anterior daughter b9.33 and the larger posterior one b9.34, although the latter lies closer to the vegetal pole.

Revisiting this division also helped clarify the developmental fates of its descendants. Early lineage-tracing in *Halocynthia* and *Ciona* (Nishida, 1987) suggested that b8.17 gives rise to both muscle and neural or endodermal strand derivatives. More recently, single-cell transcriptomic and marker gene analyses in *Ciona* refined this view by showing that b9.34 gives rise exclusively to muscle cells, whereas b9.33 produces two pairs of small “tail-tip cells” contiguous with the posterior neural tube, which likely represent a population distinct from both endodermal strand and neural cord cells (Satou et al, 2024). In *Phallusia*, the morphology, position, and size of these cells observed in our reference embryo that has reached the most advanced stage of development (Phmamm-1) support the same interpretation: the four descendants of b9.34 correspond to the “second lineage tail-muscle” (Lemaire, 2009), whereas the descendants of b9.33 form two pairs of cells at the tail tip that are clearly distinct from endodermal cells yet contiguous with the posterior nervous system, making “tail-tip cells” an appropriate designation despite their uncertain identity. This suggests that *Phallusia*, like *Ciona*, lacks a secondary endodermal strand lineage, a point that remains to be verified in *Halocynthia*. These observations also highlight the importance of tracking cells dynamically through development, a limitation that may have contributed to earlier misinterpretations of this lineage.

Taken together, the rules described in this section produce a complete, lineage-consistent extension of the Conklin naming scheme for all *Phallusia mammillata* cells up to the initial tailbud stage, including those previously unnamed or inconsistently labeled. The resulting nomenclature corrects earlier inconsistencies from the literature (e.g., b6.5, or A7.6 lineages) and aligns the *Phallusia* dataset with established descriptions in *Ciona* and *Halocynthia*.

### 2.8 Generalization of automated cell naming across individuals and species

To evaluate the generality of our automated naming strategy, we applied it to two new embryos imaged by multiview light-sheet microscopy, segmented with ASTEC (Astec, 2025; Guignard et al, 2020), and manually curated (see Methods 5.1): a *Phallusia mammillata* embryo (Phmamm-16) and an embryo of another ascidian species with transparent embryos, *Ascidiella aspersa* (Shito et al, 2023). The two genera diverged over 100 million years ago (Hasegawa et al, 2024), providing a stringent test of evolutionary conservation. Temporal alignment between *Phallusia* and *Ascidiella* embryos and the reference collection was excellent (Figure S-17), allowing one-to-one comparison of corresponding stages. In both cases, initial names at the 64-cell stage were assigned by global co-registration with the population of seven *Phallusia* reference embryos and propagated through successive divisions using local contact-area information.

Qualitative comparison with the reference embryo Phmamm-1-v2 revealed excellent positional correspondence of homologous cells in both new embryos (Figure 6a), suggesting that cell neighborhoods in the two new embryos are as canonical as those of Phmamm-1-v2, and thus that the naming accuracies of Phmamm-1-v2 (Table 4). Global geometry and local contact environments are thus conserved across more than 100 million years of ascidian evolution.

**Fig. 6:**
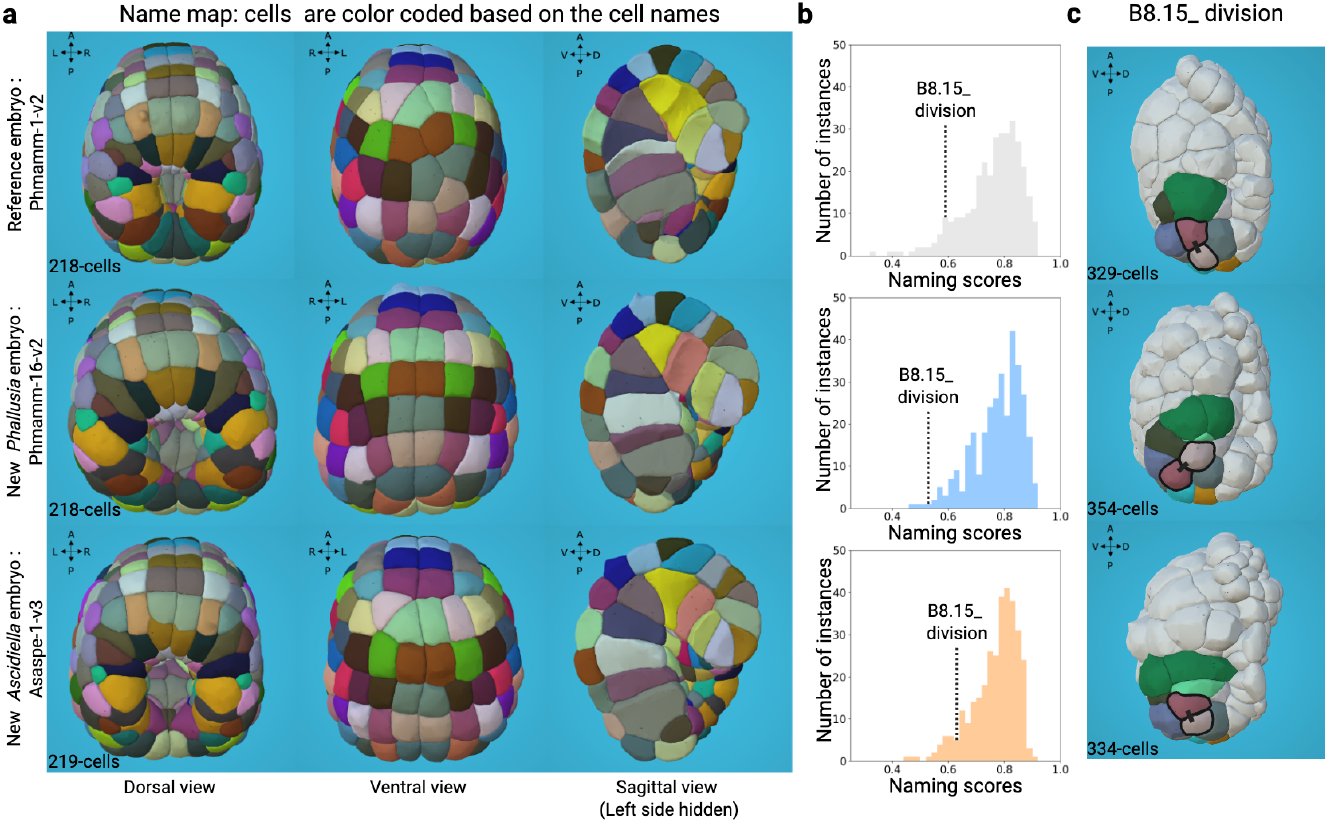
Assessment of the *de novo* naming of a new segmented embryos in *Phallusia mammillata* and *Ascidiella aspersa*. a) Dorsal (left), ventral (middle) and sagittal (right) views of a reference *Phallusia* embryo (Phmamm-1-v2, top), a new *Phallusia mammillata* embryo (Phmamm-16-v2, middle) and a new *Ascidiella aspersa* embryo (Asaspe-1-v3, bottom) at the time step corresponding to 218-cells in Phmamm-1-v2 (Figure S-17). Each pair of symmetrically matching cells is color-coded based on the cell names, allowing a qualitative comparison between Phmamm-1-v2, Phmamm-16-v2 and Asaspe-1-v3. Visualized with MorphoNet. b) Histogram of the naming scores for Phmamm-1-v2 (top, gray), Phmamm-16-v2 (middle, blue) and Asaspe-1-v3 (bottom, orange). A pairwise comparison of the naming score distributions between Phmamm-1-v2/Phmamm-16-v2 (p-value=0.148) and Phmamm-1-v2/Asaspe-1-v3 (p-value=0.288) was performed using the Kolmogorov-Smirnov (K-S) test. The position on the histograms of the B8.15 division scores are presented for Phmamm-1-v2 (naming score value=0,588), Phmamm-16-v2 (naming score value=0.533) and Asaspe-1-v3 (naming score value=0.625). c) Illustration of the atypical division angle of the B8.15 division in Phmamm-16-v2 (middle) compared to the reference embryos Phmamm-1-v2 (top) and Asaspe-1-v3 (bottom). All epidermal cells are hidden. B8.15 daughter cells are outlined in black. Black lines link B8.15 daughter cells. Common neighbor cells are color-coded according to their names. Lateral view. a,c) Antero-Posterior (AP) and Left-Right (LR) axes are represented.

To quantify naming accuracy, we used the naming score (see Supp. Methods B.7.4), which measures how typical a cell’s neighborhood configuration is relative to the reference population. High scores (close to 1) indicate canonical local topologies, whereas low scores mark atypical or ambiguous configurations. Indeed, divisions with small naming scores often reflected differences in cleavage orientation (Figure 6b,c), highlighting regions where even manual naming would have been uncertain.

This naming score confirmed high naming accuracy: the score distributions of Phmamm-16-V2, Asaspe-1-v3 and Phmamm-1-v2 were statistically indistinguishable according to the Kolmogorov–Smirnov test (Figure 6b), indicating equivalent naming reliability. At the mid-gastrula stage (218-cell stage), the naming scores for the two new embryos were overall higher than those of Phmamm-1-v2 (Figure S-18).

Taken together, these results show that our automated naming strategy generalizes robustly from one individual to another and even across species. The deep conservation of local cellular topology and embryonic geometry revealed here underscores the remarkable evolutionary stability of ascidian morphogenesis and establishes a quantitative framework for comparative ascidian embryology.

### 2.9 Naming and phenotypic analysis of experimentally manipulated *Phallusia mammillata* embryos

To test the robustness of local contact–based naming to strong morphological perturbations, we reanalyzed previously published *Phallusia mammillata* embryos in which the FGF/MEK/ERK signaling pathway, the major inducer of vegetal cell fates, had been pharmacologically inhibited (Guignard et al, 2020; Lemaire, 2009) (see Methods). These datasets display marked deviations from wild-type morpho-genesis, as MEK inhibition causes vegetal fate transformations and blocks endoderm invagination at the onset of gastrulation (Fiuza et al, 2020) (Figures 7a,c, S-20). They thus provide a stringent test of the generality of our naming framework.

**Fig. 7:**
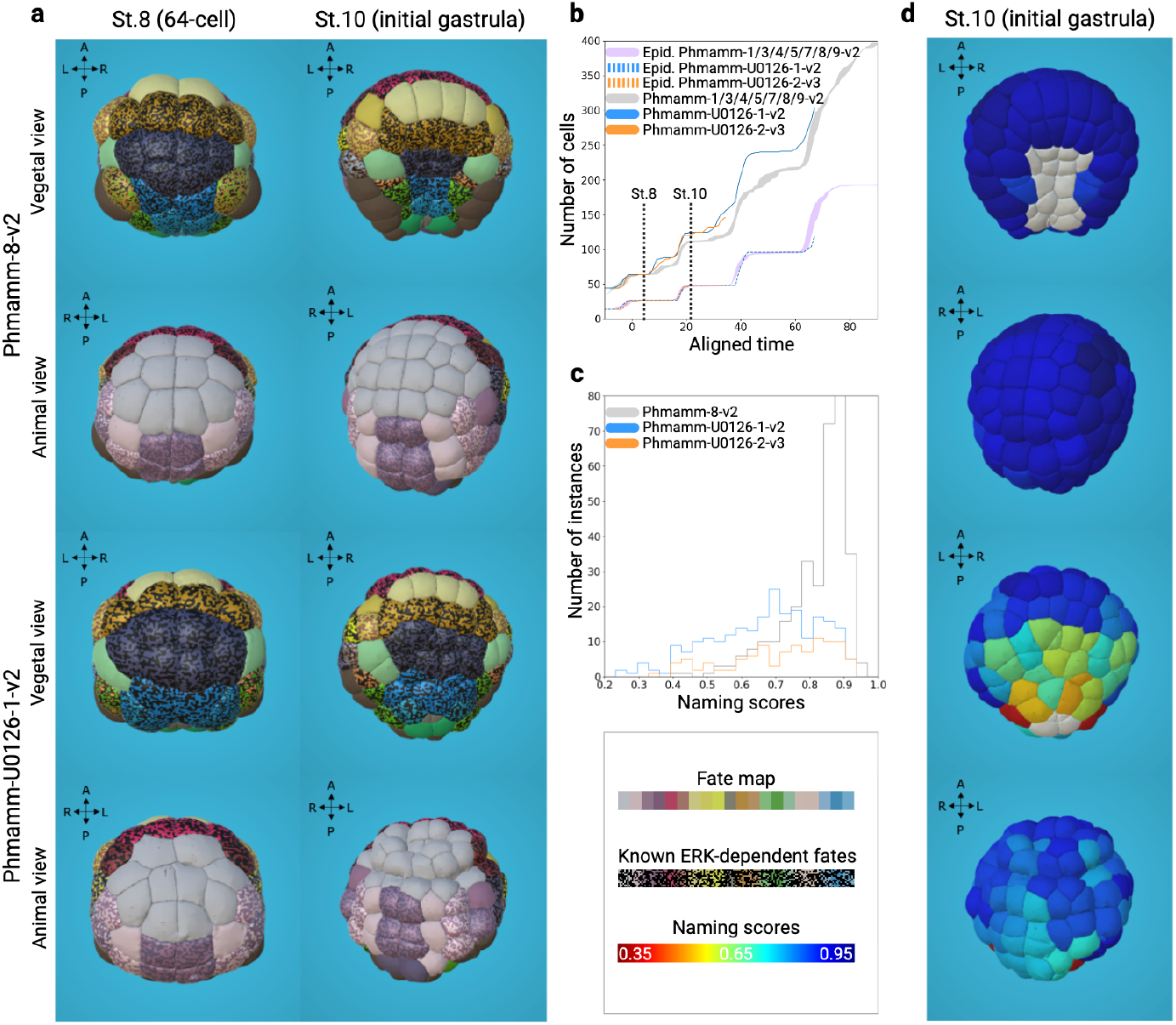
Naming and phenotypic analysis of *Phallusia* embryos treated with a MEK inhibitor from the St. 5 (16-cell). a) Vegetative and animal views of a reference embryo (Phmamm-8-v2) and an U0126-treated embryo (Phmamm-U0126-1-v2, 2µM U0126) at stages 8 (64-cell) and 10 (initial gastrula). Cells are color-coded according to the wild-type cell fates inherited from the assigned names (see Figure S-14 for the detailed tissue fates color table). In addition, multi-colored cells with black correspond to cells known to be induced by ERK signaling pathway activity or to have an ancestor that was induced by the ERK pathway earlier in development. b) Temporal alignment of reference (Phmamm-1/3/4/5/7/8/9-v2) and U0126-treated embryos based on epidermal cells. The purple curve represents the space covered by all reference embryos (Phmamm-1/3/4/5/7/8/9-v2) in their epidermal cell number evolution after temporal alignment on Phmamm-8-v2 epidermis. The dotted curves represent the evolution of epidermal cell number after temporal alignment on Phmamm-8-v2 epidermis for Phmamm-U0126-1-v2 (blue dotted line) and PhmammU0126-2-v3 (orange dotted line). The grey curve represents the space covered by all reference embryos (Phmamm-1/3/4/5/7/8/9-v2) in their whole cell number evolution after temporal alignment on Phmamm-8-v2 epidermis. The blue and orange curves represent the evolution of whole cell number after temporal alignment on Phmamm8-v2 epidermis for Phmamm-U0126-1-v2 (blue) and Phmamm-U0126-2-v3 (orange). c) Histograms of naming scores in Phmamm-8-v2 (grey), Phmamm-U0126-1-v2 (2µM U0126; blue), and Phmamm-U0126-2-v3 (6µM U0126; orange) embryos. d) Vegetative and animal views of the reference embryo Phmamm-8-v2 and of the embryo treated with U0126 (Phmamm-U0126-1-v2) at stage 10 (initial gastrula). Cells are color-coded according to the naming score. Uncolored cells in the WT embryo correspond to 7^*th*^ generation vegetal cells which have not yet divided since initialization. These cells have divided in the U0126 embryos.

Because MEK inhibition pathway respecifies vegetal cell fates, the division timings of most vegetal lineages are altered (Guignard et al, 2020), leading to differences in total cell counts between treated and wild-type (WT) embryos (Figure 7b). By contrast, the epidermal territories retain their normal identities and division patterns, providing a reliable temporal reference. Epidermal cell counts were therefore used to align developmental stages between treated and WT embryos (Figure 7b).

At the 64-cell stage, U0126-treated embryos already exhibited pronounced deviations in tissue geometry and cell topology (Figures 7a and S-19). These distortions prevented the use of the global co-registration approach applied to WT embryos. Initial cell names were therefore manually assigned, after which automated propagation across subsequent divisions was performed using local contact-area information. Despite the strong perturbation of vegetal cell divisions, this approach enabled complete, systematic naming of all cells. The distribution of naming scores, however, was shifted toward lower values compared to WT embryos (Figures 7c,d), indicating reduced confidence in cell identity assignments.

Cells of the animal hemisphere, where morphology remained largely unaffected, retained high naming scores, confirming reliable correspondence to WT homologs (Figures 7d and S-20). In contrast, vegetal cells exhibited markedly lower scores, reflecting the requirement of MEK signaling for vegetal fate specification and morphogenesis.

Together, these results demonstrate that local contact–based name propagation can robustly assign cell identities even under severe morphological perturbations, and that quantitative deviations in naming scores can serve as interpretable phenotypic readouts of developmental defects.

### 2.10 Cleavage orientation variability assessment

Having established a consistent cell naming across individuals—and demonstrated that this framework remains reliable even in the presence of strong inter-individual geometric variation—we next used it to quantify the natural variability of cell-division orientations across embryos. In earlier analyses (Results 2.6), we noted that the cleavage of cell A7.2 displayed markedly different orientations among individuals. This observation prompted a systematic assessment of how stereotyped or variable division orientations are across ascidian embryos.

To compare homologous divisions across embryos, we first defined the orientation of each division as the vector joining the barycenters of the two daughter cells immediately after cytokinesis, oriented from F(r+1).(2p) to F(r+1).(2p–1). Because left and right homologous cells divide in mirror-symmetric orientations, we reoriented all division vectors so that left–right counterparts could be directly compared. Each embryo was then aligned to a common reference at the corresponding developmental time, using homologous cell positions as landmarks. This procedure placed all division vectors into a shared spatial coordinate system, enabling quantitative cell-by-cell comparison of mean and standard deviation division orientations across embryos of the reference population (Figure 8a; see Supp. Methods B.8).

**Fig. 8:**
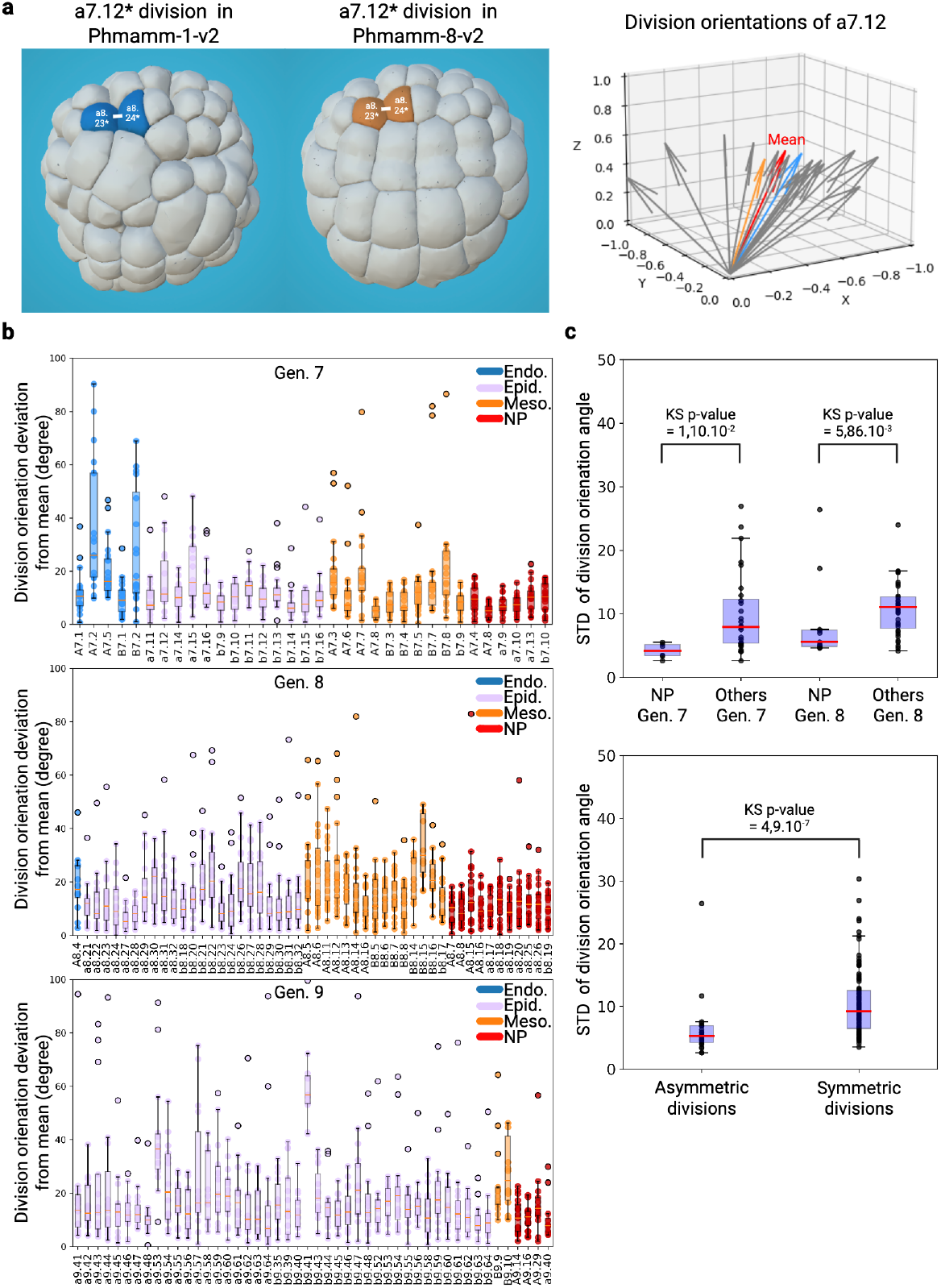
Quantification of cleavage orientation variability. a) Division orientations of cell a7.12 collected across different individuals after spatio-temporal registration. Left: division of a7.12* in Phmamm-1-v2 (blue) and Phmamm-8-v2 (orange) at the division time. Right: division orientations of a7.12 in the different individuals after spatio-temporal registration. The red vector indicates the mean division orientation of a7.12, the green vector corresponds to a7.12* division in Phmamm-1- v2, and the orange vectors to a7.12* division in Phmamm-8-v2. b) Deviation with respect to average orientation for divisions of generations 7, 8, and 9. For each cell name, the boxplot represents the deviation of its division orientations from the corresponding division mean orientation. Only divisions with at least 15 observations available (out of 20) were included. The color of the boxplot corresponds to the tissue fate of the cell. Cells contributing to multiple tissue types are duplicated, with each copy colored according to the respective tissue type. c) (Top): comparison of the standard deviation of division orientations between neural plate (NP) cells and cells from other tissues. Each scatter point corresponds to one cell name. Generation 9 was excluded due to insufficient data. NP generation 7 vs Other cells generation 7 (KS p-value = 0.011), NP generation 8 vs Other cells generation 8 (KS p-value =0.00586). (Bottom): Comparison of the standard deviation of division orientations between symmetric versus asymmetric divisions (KS p-value = 4.9*e*^−7^).

Division orientations of cells in their 7^*th*^ to 9^*th*^ generation showed widely differing levels of variability between lineages (Figure 8b). Some cells divided with highly stereotyped orientations, whereas others showed substantial deviations from the population mean. Among tissues, the neural plate displayed particularly low variability, consistent with its strong geometric organization and mechanical constraints (Hudson, 2016). Standard deviations of division orientation within the neural plate were significantly smaller than in other tissues for both the seventh and eighth mitotic cycles (Figure 8c, top). In contrast, orientation variability increased overall from generation 7 to 8 in both neural and non-neural cells, suggesting a gradual relaxation of geometric control as development progresses.

Asymmetric divisions give rise to daughter cells with distinct fates. As they often result from inductive cues from neighboring cells (Nishida, 1997), preserving these local interactions is expected to constrain their cleavage orientation. Consistent with this prediction, previously identified asymmetric divisions (Guignard et al, 2020) exhibited markedly lower orientation variability than symmetric ones (Figure 8c, bottom). A notable exception was cell B7.2, whose division orientation showed unusually high dispersion. The fate specification event in this case, however, occurs several divisions later (KB and PL, personal communication), which may lift the constraints of the division orientation of B7.2.

Among the cells with high variability in their division orientation (Figure 8b), A7.2 was previously found to exhibit distinct, recurrent orientation patterns, corresponding to alternative division modes (Figure S-16 and 9a, right). Motivated by these observations, we systematically searched for other cells exhibiting multiple preferred division orientations by clustering the orientation vectors of each cell into one or two groups and evaluating whether the resulting clusters reflected distinct division modes or merely statistical outliers (Figure 9a; see Supp. Methods B.9.2). Twenty-two cells of 7^*th*^ to 9^*th*^ generation, including A7.2, showed evidence of bimodal division orientations, implying the coexistence of alternative, recurrent division patterns within the population.

**Fig. 9:**
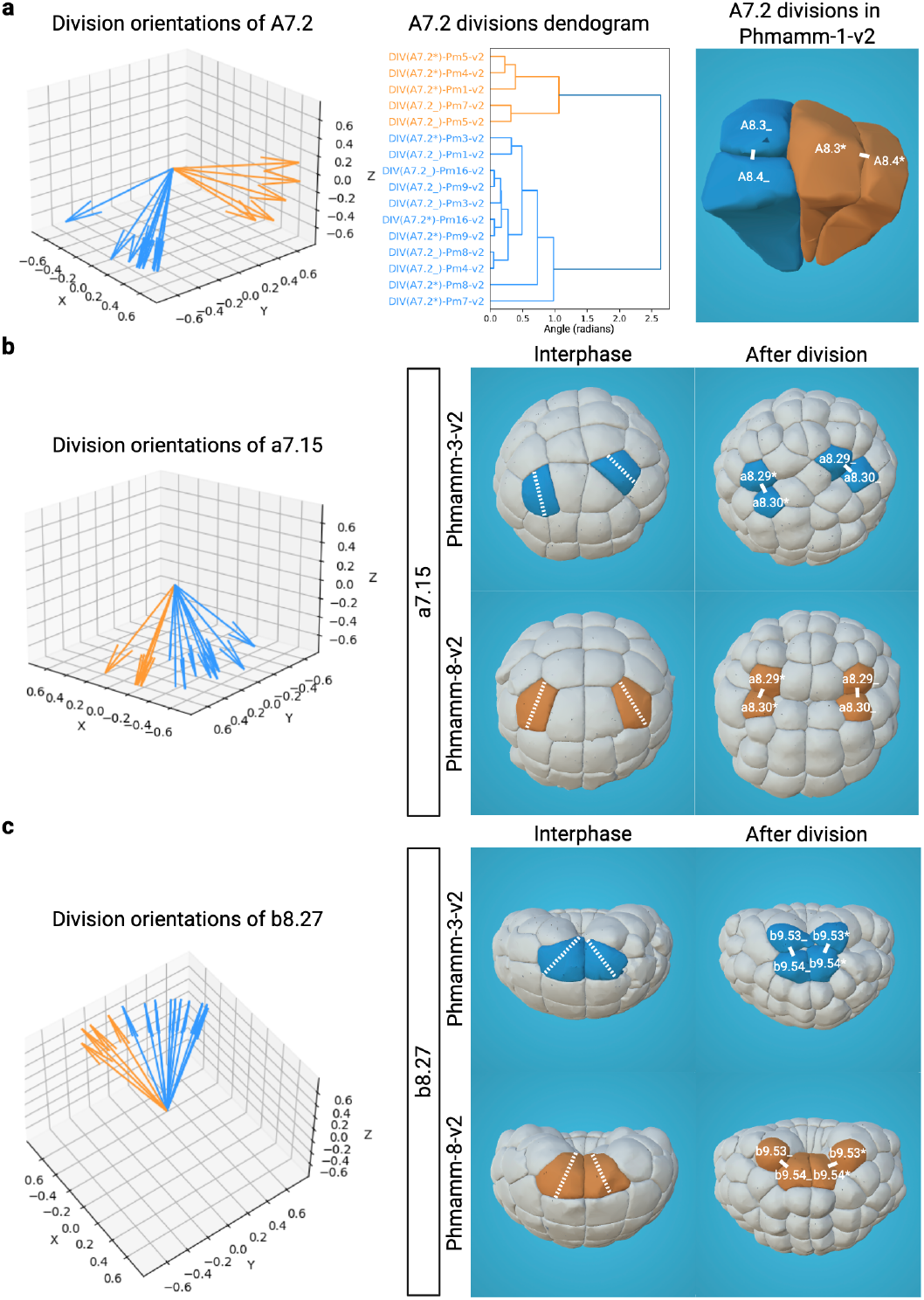
Hertwig’s rule does not explain bimodal divisions. a) Cell A7.2 with bimodal division orientation. (Left) Division orientations of A7.2 across individuals collected in the same referential clustered into two groups. (Middle) Dendrogram of A7.2 division orientations across different individuals based on the angles between orientation vectors. (Right) Division orientations of A7.2 in Phmamm-1-v2. Branches in blue correspond to division orientations clustered in blue, and similarly for orange. b) Cell a7.15 with bimodal division directions, dividing along its interphasic apical longest axis. (Left) Division orientations of a7.15 across individuals, collected in the same referential and clustered into two groups. (Right) Interphasic apical surfaces of a7.15 in Phmamm-3-v2 and Phmamm-8-v2, with the white dotted line indicating the longest apical surface axis, shown next to the corresponding divisions. c) Cell b8.27 with bimodal division directions, dividing along its interphasic apical longest axis. (Left) Division orientations of b8.27 across individuals, collected in the same referential and clustered into two groups. (Right) Interphasic apical surfaces of b8.27 in Phmamm-3-v2 and Phmamm-8-v2, with the white dotted line indicating the longest apical surface axis, shown next to the corresponding divisions.

As homologous bilateral cells could divide along distinct directions modes (Figure S-16 and 9a, right), the cell division bimodality was unlikely to be genetically predetermined and instead could arise from local mechanical or geometric factors. To explore further the impact of local geometric cues on these division patterns, we tested their conformity to Hertwig’s rule, which predicts that cells divide along their longest interphasic axis (Dumollard et al, 2017). Two representative examples are shown in Figure 9b,c: cell a7.15, whose bimodal orientations both align with its apical surface geometry, and b8.27, whose divisions violate the rule. Thus, while cell shape contributes to division orientation, additional biological cues, possibly linked to tissue-level mechanical or inductive constraints, must exist.

Taken together, this final section illustrates the power of the systematic naming framework for measuring morphogenetic precision and for dissecting how geometry, mechanics, and cell fate interact during development. Cell division orientations in ascidian embryos display contrasting patterns of variability. Most cells divide along highly reproducible orientations, particularly those undergoing asymmetric divisions associated with fate segregation. In contrast, a small subset of cells—mostly undergoing symmetric divisions—exhibit bimodal division modes, reflecting alternative yet recurrent cleavage behaviors whose underlying causes warrant further investigation.

## 3 Discussion

In this work, we developed, to our knowledge, the first framework for cell naming in an animal embryo that is automated, scalable, and tolerant to inter-individual variations. By combining global registration with local contact–based propagation, our approach achieves consistent cell naming across time, individuals, and even species, and can accommodate substantial morphological variability. This framework transforms naming from a manual, rule-based process into a quantitative operation, producing both cell identities and associated confidence scores. It therefore provides the first quantitative foundation for cell-level comparative embryology in chordates, enabling systematic cross-embryo and cross-species analyses of morphogenetic variability.

Previous automated lineage reconstructions in *C. elegans* (Bao et al, 2006) relied on canonical division patterns and deterministic lineage trees to infer cell identity. While highly effective in a species with invariant development, such approaches fail when embryos deviate from the reference lineage or exhibit local geometric variability—precisely the situations that limit their extension to other models. In contrast, our framework defines cell identity dynamically, based on local topological relationships rather than fixed division orders. This distinction allows accurate naming even when local geometry or timing diverges, as in the variable ascidian blastomeres A7.2 or B7.2, and when applied to *Ascidiella*, a species that diverged from *Phallusia* over 100 million years ago.

A central feature of our method is that it always produces a name, but with an associated naming score that quantifies how canonical a cell’s neighborhood configuration is relative to the reference population. This opens a conceptual question: how low can a score be before the assigned name loses biological relevance? In some cases, a low score may simply reflect benign geometric variation; in others, it may correspond to a true developmental divergence—for instance, a tilted division that changes which daughter receives an inductive contact, thereby altering fate specification. Because naming propagation relies on neighborhood geometry, the naming score itself can be viewed as a proxy for the preservation of inductive topology: a high score implies stable contacts and thus likely conserved signaling context, whereas a low score signals possible disruption of these cues. Future work combining cell naming with live reporters of signaling or gene expression will allow this relationship to be tested directly.

## 4 Perspectives

The framework established here constitutes the first essential building block for a multimodal atlas of ascidian development. By providing a consistent geometric and topological reference across embryos, it enables the systematic alignment of molecular, mechanical, and morphological data at single-cell resolution. Integrating transcriptomic or signaling information into this spatial scaffold will make it possible to follow, for each cell, how gene expression dynamics couple with changes in shape, position, and contact topology. Such integration will bridge molecular and physical descriptions of development, revealing how conserved geometric rules and regulatory programs jointly shape embryogenesis across individuals, conditions, and species.

## 5 Materials and methods

### 5.1 Imaging and reconstruction of new ascidian embryos from *Phallusia mammillata* and *Ascidiella aspersa*

*Phallusia mammillata* and *Ascidiella aspersa* were obtained from the Centre National de Ressources Biologiques Marines in Roscoff (France). Embryos were obtained by *in vitro* fertilisation of dechorionated eggs as in (Sardet et al, 2011). *Phallusia mammillata* and *Ascidiella aspersa* unfertilized dechorionated eggs were respectively microinjected under a stereoscope, with approximately 45pg of PH-GFP or PH-tdTomato membrane labelling synthetic mRNA as in (Guignard et al, 2020).

For lightsheet imaging, embryos were individually deposited by gravity without embedding at the bottom of a 1% Low Melt Agarose (Carl Roth, 6351.1) cone-shaped well and imaged at a constant temperature of ∼18°C in artificial seawater using a multiview lightsheet microscope (MuViSPIM, Luxendo, Germany). The PH-GFP protein was excited with a 488 nm laser (LuxX 488-60, Omicron) with simultaneous two-sided illumination. The PH-tdTomato protein was excited with a 561 nm laser with simultaneous two-sided illumination. The emitted light was collected by two opposing detection arms positioned perpendicularly to the illumination plane, resulting in simultaneous dual acquisition. Each detection arm was equipped with a 20× Olympus water dipping objective (NA 1.0) combined with a tube lens leading to a 33.3 fold image magnification. The filter used for the PH-GFP was the band-path BrightLine 525/30 filter (Semrock). The filter used for the PH-tdTomato was the long-path 561 filter. Filtered emitted light was collected by Hamamatsu V2 Flash 4 SCMOS cameras. An electronic confocal slit detection (eCSD) mode was used during image acquisition to minimise the capture of scattered light and improve contrast. The electronic slit size was matched in accordance to the diameter of the illumination beams (60 pixels). At each time point, two acquisitions were sequentially performed, the second one orthogonal to the first, generating four whole-embryo stacks, i.e. two sets of two matching 3D intensity image stacks (0, 90, 180, 270 degrees) each covering the whole imaged embryo. The embryo was imaged every two minutes with an image voxel size of 0.195 × 0.195 × 1*µm*^3^. The intensity values at each voxel were encoded in 16 bits.

The four images from each camera at a given time point were then fused to build a 3D isotropic image of the embryo using the ASTEC suite described in (Guignard et al, 2020) and freely available (Astec, 2025).

All membrane-labeled embryonic cells of the developing *Phallusia mammillata* embryo (called Phmamm-16) were then segmented at each time point using the ASTEC pipeline (Guignard et al, 2020), producing a reconstructed version 2 of the embryo integrating cell lineages and quantitative geometrical properties including cell contact surfaces, stored in an.xml file.

All membrane-labeled embryonic cells of the developing *Ascidiella aspersa* embryo (called Asaspe-1) were then segmented at each time point using the updated ASTEC pipeline (Astec, 2025), producing a reconstructed version 3 of the embryo integrating cell lineages and quantitative geometrical properties including cell contact surfaces, stored in an.xml file.

Segmentation errors (over- and under-segmentations or missing cells) were manually corrected using the MorphoNet standalone application (Gallean et al, 2025)^1^.

Finally, we applied our method to name *de novo* Phmamm-16-v2 and Asaspe1-v3. The initial naming of each cell at the 64-cell stage (St. 8) was achieved by global co-registration with our population of 7 reference embryos, and followed by the propagation of cell namings on the basis of local contact area-based information (Ascidian, 2025).

### 5.2 Collection of data from the literature

We used *Phallusia mammillata* embryos Phmamm-[1,3,4,5,7,8,9] from (Guignard et al, 2020) (MuViSPIM fusions, segmentations and embryonic properties downloaded from the project FigShare repository^2^, and explored them via MorphoNet (Gallean et al, 2025; Leggio et al, 2019)^3^. Three embryos (Phmamm-[2,6,10]) were excluded for technical or developmental reasons. We also used two previously published U0126-treated embryos (Phmamm-U0126-1 and Phmamm-U0126-2). Wild-type exclusion details, U0126 perturbation protocols and processing informations are provided in Supp. Methods B.2.

### 5.3 Dataset versioning

For clarity we annotate dataset processing versions as follows: v1 — datasets as originally published in Guignard et al (2020) (segmented with the Astec version available at publication); v2 — datasets segmented using Astec version published in (Guignard et al, 2020) and whose naming-related properties have been updated/generated using the (Ascidian, 2025) package; v3 — datasets that have been segmented with a newer (Astec, 2025) release than used in (Guignard et al, 2020) and whose naming properties were generated with (Ascidian, 2025). Detailed versioning is provided in Supp. Methods B.3.

### 5.4 Detection of manual errors affecting the Conklin syntax in the reference dataset

The “ascidian embryo” script (for more informations, see https://astec.gitlabpages.inria.fr/ascidian/ascidian_embryo.html, with the “atlas diagnosis” and “division diagnosis” parameters set to “True”, automatically parses the cell lineage and cell name information of the.xml properties file for each input.xml “atlasFiles” to check whether the cell names agrees with the Conklin syntax. This diagnosis script reports:

- Cases of inaccurate propagation of cell names through the life of a cell:
  – cell snapshots that are not named despite having a named non-dividing cell snapshot predecessor;
  – cell snapshots that are named despite having an unnamed cell snapshot predecessor;
  – cell snapshots that are named differently from their non-dividing cell snapshot predecessor;
- Cases of inaccurate propagation of cell names across a cell division:
  – cell snapshots just after a division whose name does not follow the Conklin syntax (Fr.p → [F(r+1).(2p); F(r+1).(2p-1)];
  – cell snapshots just after a division with a name compatible with the Conklin syntax, but with an unnamed sister cell;
  – cell snapshots just after a division with a name compatible with the Conklin syntax, but with a sister cell that has the same name;
  – cell snapshots just after a division with a name compatible with the Conklin syntax, but with a sister cell name that is incompatible with the Conklin syntax.

This diagnosis was launched on each Phmamm reference embryo. All detected syntax errors were corrected by manually editing the relevant.xml properties file with the sublime-text file editor. Repeating this diagnostic procedure after each round of correction ensured that no indirect syntax errors were introduced.

### 5.5 Detection and correction of manually permuted sister cell names in the reference dataset

The division distance *D*_*d*_() (see Supp. Methods B.7.2) allows to compare two divisions. We reasoned that if the names of the daughters of cell *c*_*E*_ (*i*) are inverted compared to most reference embryos *R*, the sum of the pairing distances of the permuted division of *c*_*E*_(*i*), ie (*c*_*E*_(*i*_2_), *c*_*E*_(*i*_1_)), to matching (unpermuted) divisions in reference embryos should be smaller than the sum of the pairing distances of its unpermuted division, ie (*c*_*E*_(*i*_1_), *c*_*E*_(*i*_2_)), to the reference (unpermuted) divisions. We thus constructed a Permutation Proposal Score:

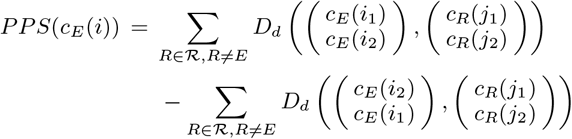

Positive values of the Permutation Proposal Score (PPS) allowed to identify sister cells whose name was likely to be permuted in the original manual naming.

As sister name permutations can affect the calculation of the PPS on neighboring sister cells and also affect the names of cells of later generations, we corrected the permuted names in the reference dataset iteratively, generation by generation, from the 7^*th*^ generation to the 10^*th*^. Starting from 7^*th*^ generation cells, we first computed the PPS for each Phmamm reference embryo using the “ascidian embryo script” (for more information, see https://astec.gitlabpages.inria.fr/ascidian/ascidian_embryo.html). We then used MorphoNet to manually inspect cells with positive PPS values in each segmented embryo. When appropriate, sister cell names were permuted by manually editing the.xml properties files of concerned Phmamm embryos. The names of the daughter cells of renamed cells were first manually deleted and then automatically renamed according to the other reference embryos using the “ascidian naming propagation” script (https://astec.gitlabpages.inria.fr/astec/ascidian_naming_propagation.html). After all 7^*th*^ generation cells were corrected, the PPS was recomputed and the following generations were iteratively analyzed in the same way.

### 5.6 Internal cell renaming

To extend Conklin’s nomenclature to internal cells, we re-evaluated the naming of sister-cell pairs belonging to internally located tissues, which had previously been assigned ad hoc without a unifying logic.

Internally located tissues were defined as all embryonic tissues except the epidermis, which remains external, and the neural plate, for which cell names were already established up to the 10^*th*^ generation (Cole and Meinertzhagen, 2004; Nicol and Meinertzhagen, 1988a,b). This analysis therefore covered all other lineages from the 8^*th*^ to the 10^*th*^ cell generations, corresponding to 98 cells.

Because cell positions evolve continuously during morphogenesis, the relative arrangement of daughter cells can change, sometimes reversing their apparent order along embryonic axes. To ensure consistent naming decisions, we therefore examined each division within a temporal window extending from one to five time points after cytokinesis. For each division, we then determined the most frequent orientation observed across the seven Phmamm reference embryos to account for inter-embryo variability.

For cells that retained contact with the exterior during gastrulation, names were assigned following Conklin’s original rule: when both daughters remained in contact with the exterior, the cell closer to the vegetal pole received the lower index; when only one did, that cell received the lower index (Conklin, 1905).

When both daughter cells were internal, we extended Conklin’s geometric logic using the MorphoNet 3D anatomical browser. Each tissue containing internal cells was visualized individually, masking all other tissues except endodermal cells, which served as a reference for the vegetal pole and the embryo’s main axes (A–P, M–L, D–V). This virtual “peeling” restored contact of internal cells with a simulated exterior, enabling manual measurement of their geodesic distance to the vegetal pole. When this distance yielded a clear ordering, the lower index was assigned to the cell closer to the vegetal pole. In ambiguous cases, we relied on the predominant division orientation across reference embryos, assigning the lower index to the more medial, anterior, or dorsal sister.

These rules were then systematically applied to all reference embryos to generate a lineage-consistent extension of the Conklin naming scheme.

### 5.7 Statistical analysis

The two-sample Kolmogorov-Smirnov (K-S) statistical analysis was performed using the scipy.stats.ks 2samp function of the scipy.stats module of the python scipy library (https://docs.scipy.org/doc/scipy/reference/generated/scipy.stats.ks_2samp.html#scipy.stats.ks_2samp). This test compares the underlying continuous distributions F(x) and G(x) of two independent samples. The null hypothesis is that the two distributions are identical, F(x)=G(x) for all x. If the KS statistic is small or the p-value is high (*>* 0.05), then we cannot reject the null hypothesis.

### 5.8 Data and code availability

Updated and newly processed datasets for this study (FigShare archives and MorphoNet direct links) are listed in Supp. Methods B.4.

Software scripts for reproducing the analyses and to perform automated de novo naming are available via astec.gitlabpages.inria.fr/ascidian/:

- “ascidian embryo” is used to assess the quality/consistency of the set of the 7 Phmamm embryos already named (see section 2.5)
- “ascidian naming timepoint” is used to name one single time point by coregistration with reference embryos (as in figure 2d-e)
- “ascidian naming propagation” is used to name divisions (sections 2.4) and to assess it with the naming score

## 6 Acknowledgements

We thank the members of the Tunicate team at CRBM for fruitful discussions throughout the course of this work and more particularly Ingrid Mormin, laboratory engineer, for essential technical and organizational support in the laboratory. We gratefully acknowledge Marc Plays and Philippe Richard’s expert animal husbandry. P.L. and G.M. were CNRS and INRIA staff scientists, respectively. K.B. was funded by an Epigenmed (ANR-10-LABX-12-01) doctoral contract and by a University of Montpellier contract extension in compensation for Covid-19 restrictions and lab closure. This work was supported by the Dig-Em (ANR-14-CE11-0013-01), Cell-whisper (ANR-19-CE13-0020) and Morphoscope (ANR-11EQPX-0029) grants of the Agence Nationale de la recherche (ANR) and by the Fondation pour la Recherche Médicale, grant number EQU202303016262, to Patrick Lemaire. We acknowledge the support and help of the Montpellier Resource Imaging (MRI) facility and in particular of Sylvain de Rossi. MRI is a core facility of the France-BioImaging infrastructure, funded by the ANR (ANR-10-INBS-04).

## Footnotes

1 https://morphonet.org/downloads

2 https://figshare.com/s/765d4361d1b073beedd5

3 https://morphonet.org/

## Notes

https://figshare.com/projects/Ascidiella_aspersa_embryonic_development/174765

https://figshare.com/projects/Phallusia_mammillata_embryonic_development/64301

### Competing Interest Statement

The authors have declared no competing interest.

