## Appendices for "Automated cell naming reveals reproducible and variable features of ascidian embryogenesis"

#### Appendix A Glossary and notations

- **Cell:** the $i^{th}$ cell of embryo $E$ is denoted $c_E(i)$. The name of cell $c_E(i)$ is denoted by $n_E(I)$
- **Cell snapshot:** thanks to the high temporal resolution of lightsheet imaging, each cell $c_E(i)$ of an embryo $E$ is imaged at several time points. A cell snapshot corresponds to $c_E(i)$ at a given time point $t$ and is denoted $c_E(i;\ t)$. All snapshots of a cell are linked by cell lineage relationships. As a consequence, quantitative information of cell snapshots $c_E(i;\ t)$ depend on time $t$.
  - $v_E(i;\ t)$ denotes the volume of cell $c_E(i)$ at time point $t$
  - $s_E(i;\ t)$ denotes the surface of cell $c_E(i)$ at time point $t$
  - $\mathbf{g}_E(i;\ t)$ denotes the centre of mass of cell $c_E(i)$ at time point $t$
- **First order neighbourhood:** neighbouring cells with which a cell $c_E(i)$ has direct physical contacts. Contacts can be transient and depend on time $t$.
  - $s_E(i,j;\ t)$ denotes the contact surface of cell $c_E(i)$ with cell $c_E(j)$ at time $t$. Cells $c_E(j)$ of the first order neighbourhood of cell $c_E(i)$ at time $t$ then verified $s_E(i,j;\ t) > 0$. By convention, we set $s_E(i,j;\ t) = 0$ for cells $c_E(j)$ that do not have a physical contact with cell $c_E(i)$.
- **Matching cells:** cells with the same Fr.p names within an embryo (bilateral cells) or across embryos (homologous cells).
- **Cell generation:** a cell generation includes all cells that are produced after the same number of divisions since the egg.
- **Primary and induced cell name differences:** a primary cell name difference corresponds to the first time that a name difference is observed in a lineage. According to Conklin's syntax, the name of a cell determines the name of its descendants, so the induced differences correspond to the observed name differences resulting from a primary cell name difference.

### Appendix B Supplemental methodology

#### B.1 Cell identification

Let us first consider how cells, prior to their naming, are identified during the segmentation process, using the ASTEC pipeline as an example (**??**). Thanks to the high temporal resolution of light-sheet imaging, each cell $c_E(i)$ of an embryo $E$ is imaged at several time points $t$. During the segmentation process, the cell snapshot $c_E(i,t)$ is identified by an arbitrary cell label, which in ASTEC has the syntax $[t][k]$, where $k$ is a unique 4-digit numerical identifier indicating the segmentation rank of the cell at time point $t$ (e.g. label 890152 indicates the 152nd cell segmented at time point 89). The different snapshots of a cell $c_E(i)$ thus receive different numerical identifiers within a segmentation, or between different segmentation runs of the same or different embryos. The pipeline yields a series of segmented images, together with a lineage that links cell snapshots at time $t$ with the ones at $t-1$ and $t+1$. After the segmentation, a number of quantitative features are computed for each cell snapshot: volume, surfaces of contact with neighboring cell snapshots, center of mass, etc. The features plus the lineage, kept in a XML file, are a representative summary of the embryo development.

#### B.2 Selection of 7 reference embryos and 2 U0126-treated embryos

We selected seven wild-type Phallusia mammillata embryos (Phmamm-1, -3, -4, -5, -7, -8, -9) from **?** as the reference population for automated naming. Three embryos were excluded: Phmamm-2 (partial exit from the field of view resulting in segmentation/tracking gaps), Phmamm-6 and Phmamm-10 (their very early pattern of cell division deviated too strongly from canonical early development for accurate manual naming of all cells).

We also included two pharmacologically perturbed datasets previously published in **?**:

- **Phmamm-U0126-1** (formerly `Astec-U0126-Pm1`): incubated with 2 $\mu$M U0126 from early stage 5a (16-cell stage). Published segmentation and XML were used as-is.
- **Phmamm-U0126-2** (formerly `Astec-U0126-Pm2`): incubated with 6 $\mu$M U0126 from the 16-cell stage. The originally published segmentation covered timepoints Tp30–Tp50; we re-segmented the fused images (Tp30–Tp79) with a newer ASTEC release (**?**) to produce `Phmamm-U0126-v3`, which was then manually curated using the MorphoNet standalone application (**?**)[1]. The curated v3 dataset is available in the project FigShare entry for Phmamm-U0126-2.

#### B.3 Dataset versioning

To ensure reproducibility and clarity we apply the following versioning scheme to all datasets:

- **v1** — Original datasets as published in **?**. Segmentations were produced with the ASTEC release used at the time of publication; lineage and naming properties correspond to the published files.
- **v2** — Datasets segmented with the ASTEC version used for the **?** publication and for which Conklin names and other naming-related properties were generated using

[1] https://morphonet.org/downloads

the ? package developed in the present study. Note that v2 entries can therefore be either datasets originally published in ? (e.g., Phmamm-1, -3, ...) or newly produced datasets that were segmented with the Astec version published in ? and named with ? (`Phmamm-16` in the present article).
- **v3** — Datasets that have been fully re-segmented using a more recent ? version than that used in ?. Example: `Phmamm-U0126-2-v1` was re-segmented to produce `Phmamm-U0126-2-v3`

#### B.4 Datasets access (FigShare & MorphoNet)

Updated and newly processed datasets for this study are provided in the project FigShare deposit (see below) and are interactively explorable on MorphoNet (see below).

- **Phmamm-1-v2**
  FigShare: https://figshare.com/account/articles/8223890
  MorphoNet: https://morphonet.org/sMObBR5f
- **Phmamm-3-v2**
  FigShare: https://figshare.com/account/articles/8235449
  MorphoNet: https://morphonet.org/T2tT2ciR
- **Phmamm-4-v2**
  FigShare: https://figshare.com/account/articles/8235455
  MorphoNet: https://morphonet.org/CQBxJ0LF
- **Phmamm-5-v2**
  FigShare: https://figshare.com/account/articles/8235458
  MorphoNet: https://morphonet.org/msz2I4pu
- **Phmamm-7-v2**
  FigShare: https://figshare.com/account/articles/8235473
  MorphoNet: https://morphonet.org/810Xf3mg
- **Phmamm-8-v2**
  FigShare: https://figshare.com/account/articles/8235479
  MorphoNet: https://morphonet.org/R6ZtvDfH
- **Phmamm-9-v2**
  FigShare: https://figshare.com/account/articles/8235482
  MorphoNet: https://morphonet.org/I36S969c
- **Phmamm-U0126-1-v2**
  FigShare: https://figshare.com/account/articles/11307431
  MorphoNet: https://morphonet.org/clBkv5B9
- **Phmamm-U0126-2-v3**
  FigShare: https://figshare.com/account/articles/11307293
  MorphoNet: https://morphonet.org/clBkv5B9
- **Phmamm-16-v2**
  FigShare: https://figshare.com/account/articles/23992869
  MorphoNet: https://morphonet.org/hdedgPfp
- **Asaspe-1-v3**
  FigShare: https://figshare.com/account/articles/23898933
  MorphoNet: https://morphonet.org/vr521gNP

#### B.5 Population normalisation

##### B.5.1 Inter-individual time normalisation

Embryos can hardly be imaged from the same exact time point of the development. Moreover, they may exhibit different development speed (Figure S-1, left). Then, for comparison purpose, we propose to synchronize the developmental time of all embryos to the one of a reference embryo $R$, with a least-squares estimation between cell count during the embryo development.

Let $N_R = \{n_{R,1}, \ldots, n_{R,i}, \ldots, n_{R,|N_R|}\}$ the different cell counts reached by embryo $R$ during the imaging period, and let $t_R(n_{R,i})$ be the acquisition time associated with $n_{R,i}$ (if there is several acquisition time with $n$ cells, $t_R(n)$ is the middle of the time interval).

To temporally register embryo $E$ onto embryo $R$, we consider the following set of cell counts

$$N = \{n \in N_R, \min(N_E) \leq n \leq \max(N_E)\} \cup \{n \in N_E, \min(N_R) \leq n \leq \max(N_R)\}$$

where duplicates have been removed. Linearly registering the developmental time of an embryo $E$ onto $R$ comes at computing

$$a_E, b_E = \arg\min_{a,b} \sum_{n \in N} \left(a t_E(n) + b - t_R(n)\right)^2$$

If there is no acquisition with $n$ cells in $R$ (resp. $E$), $t_R(n)$ (resp. $t_E(n)$) is interpolated with the acquisition times with immediate inferior and superior cell counts. Closed-form expressions for $a_E$ and $b_E$ are straightforward

$$b_E = \frac{\sum_{n \in N} t_R(n \in N) - a_E \sum_{n \in N} t_E(n)}{|N|}$$

and

$$a_E = \frac{\sum_{n \in N} t_E(n) t_R(n) - \frac{\left(\sum_{n \in N} t_E(n)\right)\left(\sum_{n \in N} t_R(n)\right)}{|N|}}{\left(\sum_{n \in N} (t_E(n))^2 - \frac{\left(\sum_{n \in N} t_E(n)\right)^2}{|N|}\right)}$$

where $|N|$ denotes the cardinal of set $N$.

##### B.5.2 Inter- and intra-individual volume normalisation

Imaged embryos exhibits a slight volume decrease during their development (Figure S-2). Therefore it is not possible to directly compare morphometric measurements (cell volume, contact surface) within an individual without compensating for this decrease. Moreover, different embryos exhibit different volumes (at the same developmental stage) and different volume decrease speed (Figure S-2).

The total volume $v_E(t)$ of an embryo $E$ exhibits a linear decrease, with variations due to imaging and/or segmentation artefacts. The linear regression is then estimated with the RANSAC[2] method which is a robust procedure. From this estimation, we get $\hat{v}_E(t) = a_E t + b_E$. This allows to get a corrective scaling factor for embryo $E$, targeting a fictitious volume $v_0$, that depends on time:

$$\alpha_E(t) = \sqrt[3]{\frac{v_0}{a_E t + b_E}}$$

[2]The python `RANSACRegressor()` method from the `scikit-learn` library.

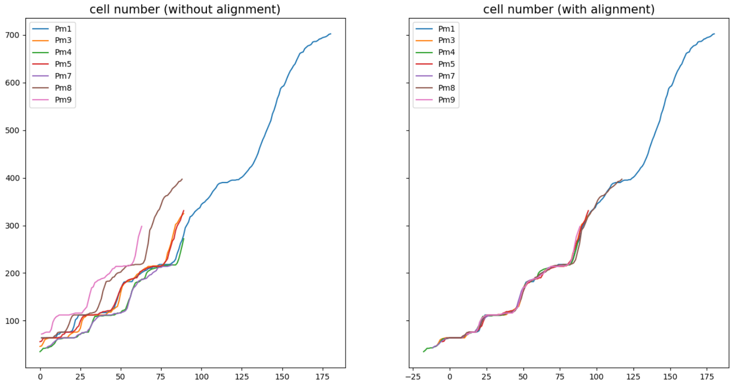


**Fig. S-1**: Left: cell counts with respect to acquisition times for the Phmamm population. Right: cell counts with respect to the synchronized acquisition times, Phmamm-1 being chosen as a reference.

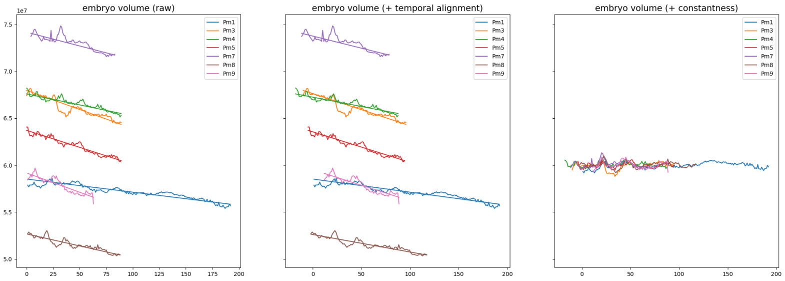


**Fig. S-2**: Embryo volume with respect to acquisition times for the Astec-Phmamm population together with estimated linear regression. Right, corrective scaling factors have been applied to get a volume of $6.10^7$ voxels.

Multiplying respectively volume and (contact) surfaces of cell snapshot $c_E(i,t)$ by $\alpha_E^3(t)$ or $\alpha_E^2(t)$ allows fair inter- and intra-individual comparisons of these measurements. Figure S-2 (right) displays the embryo volumes after volume compensation to reach a volume of $6.10^7$ voxels. Similarities of cell neighborhoods are computed with normalized contact surfaces $\tilde{s}_E(i,j,t) = \alpha_E^2(t) s_E(i,j,t)$ where $s_E(i,j,t)$ denotes the contact surface measure between cells $c(i)$ and $c(j)$ in embryo $E$ at time $t$.

When co-registering an embryo $E$ at $t_E$ onto an embryo $R$ at $t_R$, the volume difference can be compensated by the scaling matrix

$$\mathbf{S}(E(t_E), R(t_R)) \begin{pmatrix} \frac{\alpha_E(t_E)}{\alpha_R(t_R)} & 0 & 0 \\ 0 & \frac{\alpha_E(t_E)}{\alpha_R(t_R)} & 0 \\ 0 & 0 & \frac{\alpha_E(t_E)}{\alpha_R(t_R)} \end{pmatrix}$$

#### B.6 Cell naming initiation by global registration

##### B.6.1 Co-registration of embryos

Let consider two embryos, $E$ and $R$ at times $t_E$ and $t_R$, $E$ being unnamed and $R$ being named. The co-registration consists in computing the geometrical transformation that will best superimposes $E(t_E)$ and $R(t_R)$. More precisely, it will be the transformation that best superimposes the cell centres of mass (or barycenters) of $E(t_E)$ onto the ones of $R(t_R)$. For the sake of simplicity, we will drop the time notation in this section.

Since cells of $E$ have not received names, pairings between cells of $E$ and cells of $R$ are not known. Registration can then be achieved by means of the ICP (iterative closest point) iterative procedure (**?**) that alternates between 1) estimating the pairings between the two sets of points to be co-registered and 2) computing the transformation that best superimposes the built pairs. As any iterative minimization procedure, its success strongly depends on the initial conditions, here the initial relative positioning of embryos $E$ and $R$. To ensure a successful co-registration, a number of initial transformations will be tested, and the best final co-registration will be retained.

###### *Initial transformations*

The initial transformations are built by superimposing the embryo centres of mass, aligning their symmetry axis, while compensating for the embryo size difference, and adding one extra degree of freedom consisting in rotations along the symmetry axis.

The centers of mass $\mathbf{g}_E$ and $\mathbf{g}_R$ of embryos $E$ and $R$ are computed by

$$\mathbf{g}_E = \frac{1}{\sum_i v_E(i)} \sum_i v_E(i)\mathbf{g}_E(i) \quad \text{and} \quad \mathbf{g}_R = \frac{1}{\sum_i v_R(i)} \sum_i v_R(i)\mathbf{g}_R(i)$$

The unit vector $\mathbf{n}_R$ defining the (left-to-right) symmetry axis of the named embryo $R$ is computed by normalizing the vector

$$\frac{1}{\sum_{i \in N_*(R)} v_R(i)} \sum_{i \in N_*(R)} v_R(i)\mathbf{g}_R(i) - \frac{1}{\sum_{i \in N\_(R)} v_R(i)} \sum_{i \in N\_(R)} v_R(i)\mathbf{g}_R(i)$$

where $N_*(R)$ and $N\_(R)$ denotes respectively the sets of right cells (whose names are ending by '$*$') and left cells (whose names are ending by '_').

The unit vector $\mathbf{n}_E$ defining the symmetry axis of the unnamed embryo $E$ is computed as in (**?**). However, it can be either the left-to-right or the right-to-left vector, thus we will assess the two vectors $\varepsilon\mathbf{n}_E$ with $\varepsilon \in \{-1, 1\}$.

Initial transformations $\mathbf{A}(\varepsilon, k)$ are finally computed by

$$\mathbf{A}(\varepsilon, k) = \mathbf{R}(\varepsilon\mathbf{n}_E(k), \mathbf{n}_R)\mathbf{S}(E, R)\mathbf{R}(\mathbf{n}_R, \theta_k)$$

where

- $\mathbf{R}(\varepsilon\mathbf{n}_E(k), \mathbf{n}_R)$ is a rotation matrix that aligns $\mathbf{n}_R$ onto $\varepsilon\mathbf{n}_E(k)$,
- $\mathbf{S}(E, R)$ is a scaling matrix that compensates for embryo size differences (Supp. Methods B.5.2), and
- $\mathbf{R}(\mathbf{n}_R, \theta_k)$ is a rotation of axis $\mathbf{n}_R$ and angle $\theta_k$, where the set $\{\theta_k = k\delta\theta\}$ is a sampling of $[0, 2\pi[$ ($\delta\theta$ being set to 15 degrees).

##### *Registration*

Each initial transformation $\mathbf{A}(\varepsilon, k)$ is refined through an ICP (iterative closest point) procedure (?) that registers the set of cell centres of mass $\{\mathbf{g}_E(i)\}$ of embryo $E$ onto the set of cell centres of mass $\{\mathbf{g}_R(j)\}$ of embryo $R$.

Our implementation of ICP iteratively built pairings between $\{\mathbf{g}_E(i)\}$ and $\{\mathbf{g}_R(j)\}$ and estimated an affine transformation from the pairings with a least-trimmed-squares procedure (?) to account for possible outliers (which may arise when the embryo cell compositions are different): these two steps are iterating until convergence, yielding a refined transformation $\hat{\mathbf{A}}(\varepsilon, k)$ for each initial transformation $\mathbf{A}(\varepsilon, k)$.

Let $cc_R(\mathbf{g})$ denotes the point of $\{\mathbf{g}_R(j)\}$ that is closest to point $\mathbf{g}$

$$cc_R(\mathbf{g}) = \mathbf{g}_R \left( \arg\min_j \|\mathbf{g} - \mathbf{g}_R(j)\| \right)$$

The transformation that registers $\{\mathbf{g}_E(i)\}$ onto $\{\mathbf{g}_R(j)\}$ can then be assessed by the sum of residuals, where each point $\mathbf{g}_E(i)$ is paired with its closest point in $\{\mathbf{g}_R(j)\}$ after transformation:

$$SR(\hat{\mathbf{A}}(\varepsilon, k)) = \frac{1}{|\{\mathbf{g}_E(i)\}|} \sum_i \left| \hat{\mathbf{A}}(\varepsilon, k)\mathbf{g}_E(i) - cc_R\left(\hat{\mathbf{A}}(\varepsilon, k)\mathbf{g}_E(i)\right) \right|$$

where $|\{\mathbf{g}_E(i)\}|$ denotes the cardinal of $\{\mathbf{g}_E(i)\}$. To account for outliers, we discard the 20% of highest residuals, as in (?), and consider as assessment score the average of the 80% smallest residuals, that we denote by $SR_{80}()$.

We retain as the transformation $\hat{\mathbf{A}}_{R\leftarrow E}$ to superimpose the embryo $E$ onto $R$ the one that minimize $SR_{80}()$:

$$\hat{\mathbf{A}}_{R\leftarrow E} = \arg\min_{\varepsilon} \min_{k} SR_{80}(\hat{\mathbf{A}}(\varepsilon, k))$$

##### B.6.2 Transfer of cell names between aligned embryos

We establish pairs of reciprocally closest cell snapshot centres of mass between $\{\mathbf{g}_E(i)\}$ and $\{\mathbf{g}_R(j)\}$ after transformation $\hat{\mathbf{A}}_{E\leftarrow R}$, which are defined by

$$\left\{ \left( \mathbf{g}_E(i), \mathbf{g}_R(j_R(i)) = cc_R(\hat{\mathbf{A}}_{R\leftarrow E}\mathbf{g}_E(i)) \right) \text{ such that } cc_E(cc_R(\hat{\mathbf{A}}_{R\leftarrow E}\mathbf{g}_E(i))) = \mathbf{g}_E(i) \right\}$$

Note that there may be cell snapshot centres of mass from embryo $E$ that can remain unpaired. We assume that paired cells are likely to have the name, then that cell $c_E(i)$ is likely to be named as $c_R(j_R(i))$, ie $n_R(j_R(i))$.

If we add a number of named embryos $R_k$ at hand, each cell $c_E(i)$ of $E$ will have a set of candidate names $CN_E(i) = \{n_{R_k}(j_{R_k}(i))\}$ issued from all $R_k$. We design an ad-hoc voting procedure to attribute one single name to each cell $c_E(i)$ (Algorithm 1). Again, it may happen that some cells remain unnamed.

#### B.7 Distances from contact surfaces

We estimated the distance between cells, or between divisions, from the neighborhood information: we hypothesized that the same cell (of given name) has similar neighborhood across a population of embryos.

**Algorithm 1** Naming from sets of candidate names

**Require:** $CN_E(i)$ for cells $c_E(i)$
**repeat**
  $AttributedNames = \emptyset$
  **for** sets $CN_E(i)$ that contains only occurrences of one single name **do**
    Give that name of $c_E(i)$
    Add that name to $AttributedNames$
  **end for**
  Remove names of $AttributedNames$ from remaining sets $CN_E(i)$
**until** no changes occurs

##### B.7.1 Neighborhood distance

Let consider two cell snapshots, $c_E(i;\ t_E)$ and $c_R(i;\ t_R)$, respectively extracted from embryo $E$ at time $t_E$ and embryo $R$ at time $t_R$. Each cell can be represented by the vector of its contact surfaces (after volume normalization), $c_E(i;\ t_E) \triangleq (s_E(i,k;\ t_E))_k$. A distance between two cells can be built by comparing the vectors of contact surfaces, where contact surfaces are paired according to the neighbor cell name.

Let $N_E(i;\ t_E) = \{k / s_E(i,k;\ t_E) \neq 0\}$ be the set of neighbors of cell snapshot $c_E(i;\ t_E)$, a Manhattan distance between the contact surface vectors, where neighbors of same name are compared, defines the distance $|c_E(i;\ t_E) - c_R(i;\ t_R)|_S$,

$$|c_E(i;\ t_E) - c_R(i;\ t_R)|_S \triangleq \sum_{(k,\ell), n_E(k;\ t_E) = n_R(\ell;\ t_R)} |s_E(i,k;\ t_E) - s_R(j,\ell;\ t_R)|$$

with the convention that the contact surfaces of non-neighboring cells are set to zero: $s_E(i,k;\ t_E) = 0$ if $k \notin N_E(i;\ t_E)$.

Such a distance can assess the proximity of two cells when the embryo cell compositions are the same (eg at 64-, 76-, or 112- cells stages). It will however be biased by the division heterochrony at any other time points (Figure S-12). Typically, a mother cell (eg A7.2*) belongs to one neighborhood (say the one of $c_E(i;\ t_E)$) while the daughter cells (here A8.3* and A8.4*) belongs to the other neighborhood (here the one of $c_R(i;\ t_R)$). We then fused neighbors (which comes to add surface contacts) to build a virtual parent cell, so the cell neighborhoods can be faithfully compared. These virtual neighborhoods are made of the closest common ancestors between the neighborhoods to be compared. It comes to generalize the above distance with paired sets of neighbors

$$|c_E(i;\ t_E) - c_R(j;\ t_R)|_S \triangleq \sum_{(K,L)} \left| \sum_{k \in K} s_E(i,k;\ t_E) - \sum_{\ell \in L} s_R(j,\ell;\ t_R) \right|$$

When neighbor fusion occurred, one of the two paired sets is a singleton (the mother cell) while the other set contains the daughter cells.

While providing an absolute measure of the proximity of two cells, the distance $|c_E(i;\ t_E) - c_R(i;\ t_R)|_S$ does not allow to faithfully compare the distances between any couple of cells since they depend of the cell surface. We prefer introduce a relative measure of the proximity of two cells, named neighborhood distance (although it is not a distance in a mathematical sense) denoted $D_n()$, which is the above Manhattan distance divided by the sum of the two cell surfaces, $s_E(i;\ t_E)$ and $s_R(j;\ t_R)$. It can be interpreted as the fraction of the cell surfaces that contact different neighbours.

$$D_n(c_E(i;\ t_E) - c_R(j;\ t_R)) \triangleq \frac{|c_E(i;\ t_E) - c_R(j;\ t_R)|_S}{s_E(i;\ t_E) + s_R(j;\ t_R)}$$

Since $s_E(i;\ t_E) = \sum_k s_E(i,k;\ t_E)$ and $s_R(j;\ t_R) = \sum_\ell s_R(j,\ell;\ t_E)$, we have $D_n(c_E(i;\ t_E) - c_R(j;\ t_R)) \in [0,1]$ with

- $D_n(c_E(i;\ t_E) - c_R(j;\ t_R)) = 0$ the two cells have exactly the same contact surfaces with the same cells, and
- $D_n(c_E(i;\ t_E) - c_R(j;\ t_R)) = 1$ the two cells do not have any common contact surfaces.

##### B.7.2 Division distance

A division is represented by the ordered daughter cell snapshots, eg $d_E(i) \triangleq \begin{pmatrix} c_E(i_1;\ t) \\ c_E(i_2;\ t) \end{pmatrix}$ where $c_E(i_1)$ and $c_E(i_2)$ are the cells resulting from the division of cell $c_E(i)$. According that division of cell $c_E(i)$ occurs at time $td_E(i)$, and thanks to the stability of the local neighborhood (Results **??**), the division of $c_E(i)$ can be represented by its two daughter cell snapshots at $T_E(i) = td_E(i) + 4\delta t$ (the fourth time point after the division)

$$d_E(i) \triangleq \begin{pmatrix} c_E(i_1;\ T_E(i)) \\ c_E(i_2;\ T_E(i)) \end{pmatrix}$$

The distance between two divisions is based upon the neighborhood distances of the paired daughter cells. As for the neighborhood distance, a normalization by the sum of the four daughter cell surfaces ensures to get a measure in $[0,1]$. The division distance, denoted $D_d()$, is defined by

$$\begin{aligned} D_d(d_E(i), d_R(j)) &= D_d\left( \begin{pmatrix} c_E(i_1;\ T_E(i)) \\ c_E(i_2;\ T_E(i)) \end{pmatrix}, \begin{pmatrix} c_R(j_1;\ T_R(j)) \\ c_R(j_2;\ T_R(j)) \end{pmatrix} \right) \\ &\triangleq \frac{|c_E(i_1;\ T_E(i)) - c_R(j_1;\ T_R(j))|_S + |c_E(i_2;\ T_R(i)) - c_R(j_2;\ T_R(j))|_S}{s_E(i_1;\ T_E(i)) + s_E(i_2;\ T_E(i)) + s_R(j_1;\ T_R(j)) + s_R(j_2;\ T_R(j))} \end{aligned}$$

For the sake of simplicity, we will drop out the time from the division distance from now on.

##### B.7.3 Naming a division with respect to a collection of reference embryos

Let $\{R_k\}$ be a collection of embryos, with named divisions. By convention, the upper cell in division $d_R(j) = \begin{pmatrix} c_R(j_1) \\ c_R(j_2) \end{pmatrix}$ is the one with the odd position integer (F(r+1).(2p-1)) while the lower one is the one with the even position integer (F(r+1).(2p)).

Naming a division $d_E(i)$ comes to decide the daughter cell order, ie whether the division $d_E(i)$ should be written $\begin{pmatrix} c_E(i_1) \\ c_E(i_2) \end{pmatrix}$ or $\begin{pmatrix} c_E(i_2) \\ c_E(i_1) \end{pmatrix}$.

We then compared the similarities with respect to the reference embryos $R_k$, and named $d_E(i)$ with the choice that yields a minimum of average similarity. For instance, if $\sum_k D_d\left( \begin{pmatrix} c_E(i_1) \\ c_E(i_2) \end{pmatrix}, \begin{pmatrix} c_{R_k}(j_1) \\ c_{R_k}(j_2) \end{pmatrix} \right) < \sum_k D_d\left( \begin{pmatrix} c_E(i_2) \\ c_E(i_1) \end{pmatrix}, \begin{pmatrix} c_{R_k}(j_1) \\ c_{R_k}(j_2) \end{pmatrix} \right)$,

$c_E(i_1)$ and $c_E(i_2)$ are respectively as $c_R(j_1)$ and $c_R(j_2)$.

This is similar to the Permutation Proposal Score (Methods **??**), except that the division $c_E(i)$ is not yet named.

##### B.7.4 Naming score

We define a naming score that assesses the likelihood to find similar division patterns in the population of reference embryos.

The division distance $D_d()$ (Supp. Methods B.7.2) allows to compare two divisions, providing a straightforward means to assess the named divisions. Using all the reference embryos at hand may provide a non-robust assessment since some reference cleavage orientations may largely differ from the evaluated one, as it is the case for divisions exhibiting different cleavage orientation pattern (as A7.2, see Figure S-16).

To assess whether similar division patterns exist in the reference population, the naming score only uses the *closest* (the ones with the smallest distance) exemplars in the population, allowing to discard the most dissimilar division patterns:

$$NS(c_E(i)) = 1 - \sum_{R \in \mathcal{R}(c_E(i))} D_d\left( \begin{pmatrix} c_E(i_1) \\ c_E(i_2) \end{pmatrix}, \begin{pmatrix} c_R(j_1) \\ c_R(j_2) \end{pmatrix} \right)$$

where $\mathcal{R}(c_E(i))$ denotes this set of closest exemplars. To assess the named divisions, we thus used the closest two reference embryos.

This naming score lies in [0, 1], and large values indicates that similar division neighborhoods can be found in the population of reference embryos, while small values indicates a non-typical neighbourhood.

#### B.8 Mean orientation computation

For a set of orientations represented as points on the unit sphere, the mean division direction was computed using an iterative tangential projection method on the sphere.

Let $\{\mathbf{u_k}\}_{k=1}^{K}$ be unit vectors on the sphere $S^2 \subset \mathbb{R}^3$. A distance on the sphere is the arc length, i.e. an angle (in radians) on the unit sphere. The distance between the two vectors $\mathbf{x}, \mathbf{y} \in S^2$ is then given by $d(\mathbf{x}, \mathbf{y}) = \arccos(\mathbf{x} \cdot \mathbf{y})$.

The Fréchet mean (or Riemannian mean) is the unit vector $\hat{\mathbf{n}} \in S^2$ minimizing the sum of squared geodesic distances: $\hat{\mathbf{n}} = \arg\min_{\mathbf{n} \in S^2} \sum_{k=1}^{K} d(\mathbf{n}, \mathbf{u_k})^2$.

There is no close form to compute this expression, but an iterative scheme can calculate $\hat{\mathbf{n}}$ as follows:

1. Initialize $\mathbf{n}_0$ as either one of the $\mathbf{u}_k$ or the normalized Euclidean mean $\frac{\sum_k \mathbf{u}_k}{\|\sum_k \mathbf{u}_k\|}$.
2. At iteration $i$, for each $\mathbf{u}_k$, define the projection on the plane tangent to the sphere on $\mathbf{n}_i$: $\mathbf{p}_k = \mathbf{n}_i + \theta_k \frac{\mathbf{u}_k - (\mathbf{n}_i \cdot \mathbf{u}_k)\mathbf{n}_i}{\|\mathbf{u}_k - (\mathbf{n}_i \cdot \mathbf{u}_k)\mathbf{n}_i\|}, \quad \theta_k = d(\mathbf{n}_i, \mathbf{u}_k)$.
3. Compute the barycenter of $\mathbf{p}_k$'s on the tangent plane:
   $\mathbf{q}_{i+1} = \mathbf{n}_i + \frac{1}{K} \sum_{k=1}^{K} \theta_k \frac{\mathbf{u}_k - (\mathbf{n}_i \cdot \mathbf{u}_k)\mathbf{n}_i}{\|\mathbf{u}_k - (\mathbf{n}_i \cdot \mathbf{u}_k)\mathbf{n}_i\|}$.
4. Project back this barycenter onto the sphere:
   define $\alpha_i = \|\mathbf{q}_{i+1} - \mathbf{n}_i\|, \qquad \mathbf{r}_i = \frac{\mathbf{n}_i \times \mathbf{q}_{i+1}}{\|\mathbf{n}_i \times \mathbf{q}_{i+1}\|}$,
   and update using Rodrigues' formula:
   $\mathbf{n}_{i+1} = \mathbf{n}_i + \sin(\alpha_i)(\mathbf{r}_i \times \mathbf{n}_i) + (1 - \cos(\alpha_i))\left(\mathbf{r}_i \times (\mathbf{r}_i \times \mathbf{n}_i)\right)$.

Repeat until convergence; the limit point is the mean orientation $\hat{\mathbf{n}}$.

#### B.9 Modes of cell division orientation distribution

##### B.9.1 1D illustrative example

Suppose we have a set of samples drawn from a one-dimensional random variable $x \in \mathbb{R}^1$. From these observations, we can estimate the distribution of $x$ using kernel density estimation (KDE) (**?**), where each sample is assigned a kernel (here a Gaussian) with width $\sigma$. The choice of $\sigma$ strongly affects the number of modes: a sufficiently large $\sigma$ will yield a unimodal distribution, while a sufficiently small $\sigma$ (approaching a Dirac kernel) yields as many modes as samples.

Several general rules have been proposed to estimate an appropriate bandwidth (e.g., Student's rule). However, these methods assume a large number of samples. In our case, the number of observations is limited, so we restrict our analysis to detect at most two possible modes (Figure S-3).

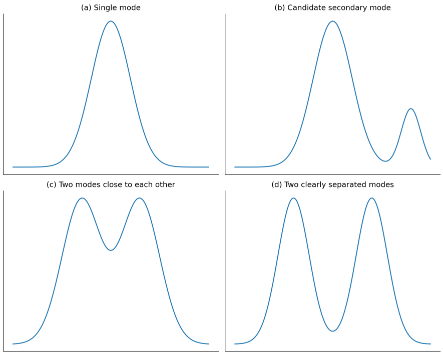


**Fig. S-3**: We distinguish between four cases: (a) a single mode, (b) a candidate secondary mode, so the number of samples contributing the second mode is small hence might be due to outliers (will need a larger number of observations to conclude) (c) two modes close to each other (close distance between modes), and (d) two clearly separated modes (with a large distance between them and a comparable number of samples contributing to each mode).

##### B.9.2 Application to division orientations

We propose a framework to detect bimodal division orientations (Figure S-3). For a given division $c_i$, we define the set $\mathcal{V}_i$ of orientation vectors across embryos $\{E^e\}$ (collected in the same referential):

$$\mathcal{V}_i = \{\mathbf{v}_i^e \mid e \in \{E^e\}\},$$

where the unit vector $\mathbf{v}_i^e \in \mathbb{R}^3$ represents also a point on the unit sphere.

The distribution of division orientations $\mathcal{V}_i$ was estimated using a Gaussian kernel density estimator on the sphere (**?**):

$$\mathrm{PDF}(\mathbf{n}) = \frac{1}{\mathsf{V}_i} \sum_{e \in E^e} K_\sigma(\mathbf{n}, \mathbf{v}_i^e),$$

with

$$K_\sigma(\mathbf{n}, \mathbf{v}_i^e) = \frac{1}{C} \exp\left( -\frac{\left(\arccos(\mathbf{n} \cdot \mathbf{v}_i^e)\right)^2}{2\sigma^2} \right),$$

where $C$ is the normalization constant and the angular distance defines the metric. This yields a probability distribution over the unit sphere, where each voxel corresponds to the likelihood of observing a division orientation at that direction.

Because of the limited number of observations per division, we hypothesized that each cell could exhibit at most two distinct orientation patterns. The kernel width $\sigma$ was progressively increased (starting at 0.05 radians and incremented by 0.1) until two modes appeared. Every voxel (and each observation) was then assigned to a mode using a reverse watershed algorithm, clustering the division orientations into two groups.

We evaluated each division orientations clustering to determine whether the identified clusters represented truly distinct groups (bimodal divisions) or instead reflected a single underlying pattern (stereotyped divisions). This assessment was based on the following criteria:

- **Isolation**: the angular separation between the two cluster modes, $\theta_m = \arccos(\mathbf{m}_1 \cdot \mathbf{m}_2)$, where $\mathbf{m}_1$ and $\mathbf{m}_2$ denote the identified modes.
- **Balance**: whether both clusters contained a sufficient number of observations to be biologically meaningful, rather than being driven by a few outliers, $p = \max\left(\frac{N_1}{N_1+N_2}, \frac{N_2}{N_1+N_2}\right)$, where $N_k$ is the number of vectors in cluster $k$. A value of $p = 0.5$ corresponds to equally sized clusters, while $p$ close to 1 indicates strong imbalance, suggesting that the secondary mode may be inconclusive.

Figures S-4, S-5, and S-6 summarize this analysis across generations. High isolation (higher than 40 degrees) may indicate divisions with two distinct orientation patterns, while high imbalance (higher than 0.9) points to inconclusive cases due to possible outliers. Divisions requiring relatively large kernel widths to get two modes distributions are particularly noteworthy, as they may reflect high intra-cluster variability, possible outliers, or the presence of a third pattern.

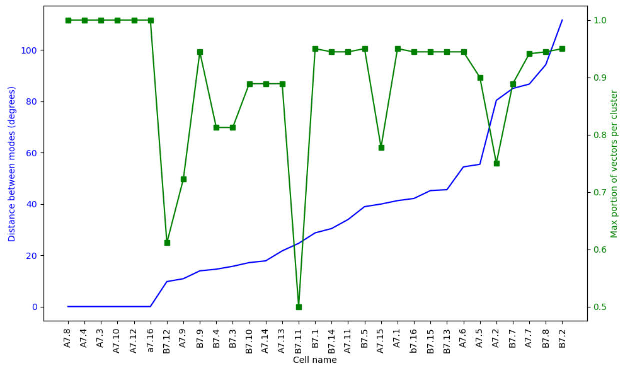


**Fig. S-4**: Clustering analysis of division orientations at the $7^{th}$ generation. The blue curve (left $y$-axis) shows the isolation between the two cluster means. The green curve (right $y$-axis) indicates the maximum cluster proportion, a measure of balance.

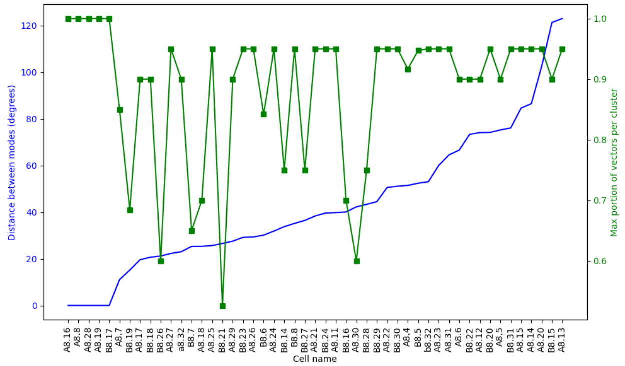


**Fig. S-5**: Clustering analysis of division orientations at the $8^{th}$ generation. The blue curve (left $y$-axis) shows the isolation between the two cluster means. The green curve (right $y$-axis) indicates the maximum cluster proportion, a measure of balance.

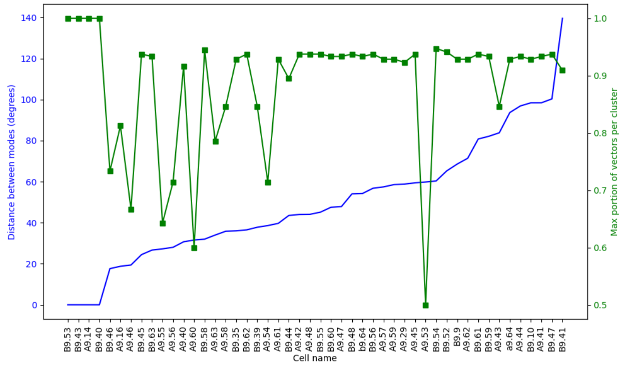


**Fig. S-6**: Clustering analysis of division orientations at the $9^{th}$ generation. The blue curve (left $y$-axis) shows the isolation between the two cluster means. The green curve (right $y$-axis) indicates the maximum cluster proportion, a measure of balance.

In the $7^{th}$ generation, we identified cells such as B7.2 (and A7.2, B7.7, A7.7, B7.8) that displayed high intra-cluster variation (due to high kernel width) (Figure S-7).

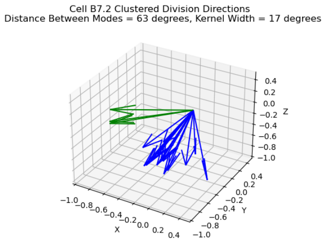


**Fig. S-7**: Clustered division orientations of B7.2 from the population showing two patterns.

#### B.10 Intra- versus inter-individual variation

We assessed intra- versus inter-individual variation, for all stereotyped divisions we compared the distribution of the angles between division orientations in each embryo $E^e$ and their corresponding symmetrical counterparts $E^{sym-e}$:

$$\theta_{e,sym-e}(Fr.p) = \arccos\left(\mathbf{v}^{e}_{Fr.p} \cdot \mathbf{v}^{sym-e}_{Fr.p}\right),$$

and the angles distribution between division orientations of homologous cells in $E^e$ to $\{E^f\}_{f \notin \{e,sym-e\}}$:

$$\theta_{e,f}(Fr.p) = \arccos\left(\mathbf{v}^{e}_{Fr.p} \cdot \mathbf{v}^{f}_{Fr.p}\right), \quad f \notin \{e, sym-e\}.$$

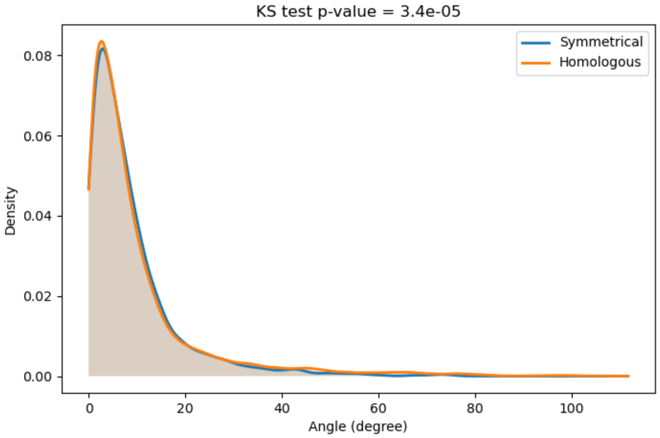


**Fig. S-8**: Distributions of cell division orientation angles $\{\theta_{e,sym-e}(Fr.p)\}$ and $\{\theta_{e,f}(Fr.p)\}$ for the $7^{th}$ generation cells.

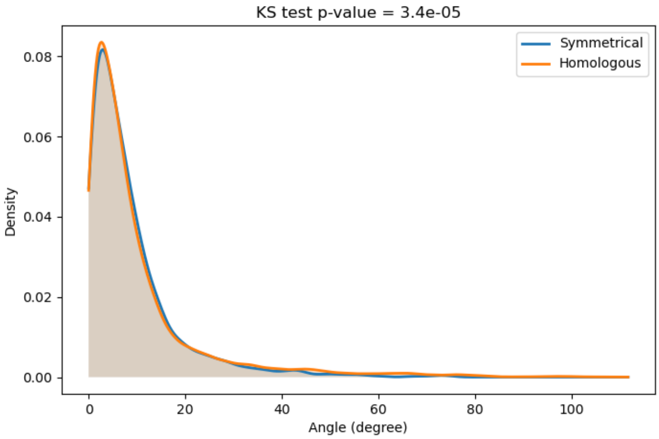


**Fig. S-9**: Distributions of cell division orientation angles $\{\theta_{e,sym-e}(Fr.p)\}$ and $\{\theta_{e,f}(Fr.p)\}$ for the $8^{th}$ generation cells.

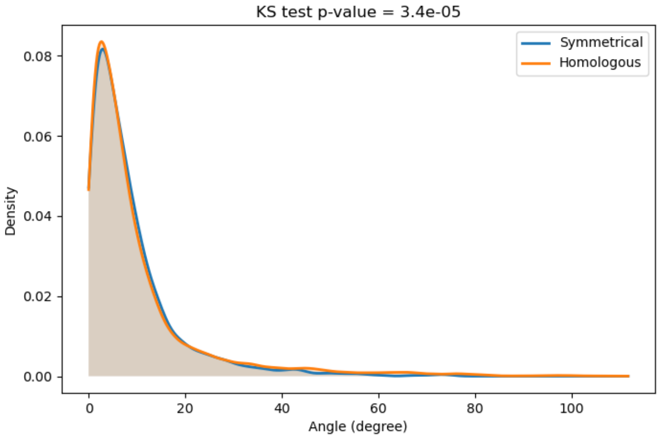


**Fig. S-10**: Distributions of cell division orientation angles $\{\theta_{e,sym-e}(Fr.p)\}$ and $\{\theta_{e,f}(Fr.p)\}$ for the $9^{th}$ generation cells.

The reference embryo $r$ used for spatio-temporal alignment, was chosen iteratively as each embryo in turn to avoid bias in reference selection. The distributions were estimated by KDE using a Gaussian kernel, with the bandwidth selected according to Scott's rule. The distributions for each generation of $\{\theta_{e,sym-e}(Fr.p)\}$ and $\{\theta_{e,f}(Fr.p)\}$ were similar. This analysis shows that intra- and inter-embryo variation for stereotyped divisions is similar.

### Appendix C Supplemental information

| Embryo ID | # misnamed cells due to manual errors in the naming syntax (*) | # PPS | # PPS accepted | # renamed cells following PPS analysis (*) |
|---|---|---|---|---|
| Phmamm-1-v1 | 9 | 10 | 10 | 24 |
| Phmamm-3-v1 | 2 | 9 | 9 | 22 |
| Phmamm-4-v1 | 2 | 3 | 3 | 6 |
| Phmamm-5-v1 | 2 | 17 | 17 | 44 |
| Phmamm-7-v1 | 0 | 1 | 1 | 2 |
| Phmamm-8-v1 | 19 | 7 | 7 | 38 |
| Phmamm-9-v1 | 2 | 7 | 7 | 14 |
| Total | 36 | 54 | 54 | 150 |

**Table C1**: **Summary of the manual errors corrected using the Permutation Proposal score.** (#)="Number of". (*): Includes both primary and induced errors (see glossary A).

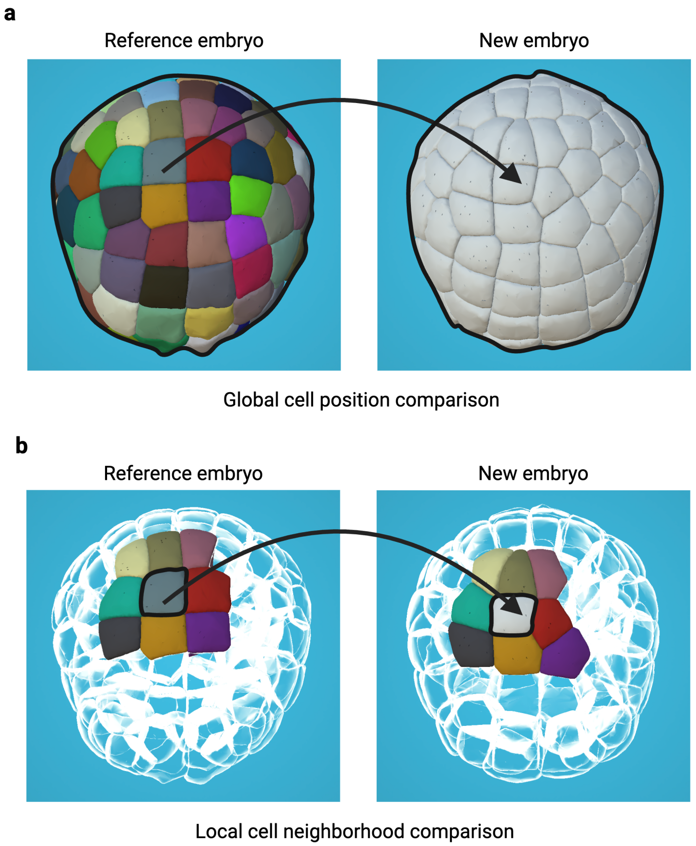


**Fig. S-11**: **Two different naming strategies.** a) Naming by global cell position comparison. b) Naming by local cell neighborhood comparison. a-b) Named cells are shown in color. Unnamed cells have no color. Cells not used for naming are transparent. The cells surrounded in black indicate the cells for which names are transferred by the strategy.

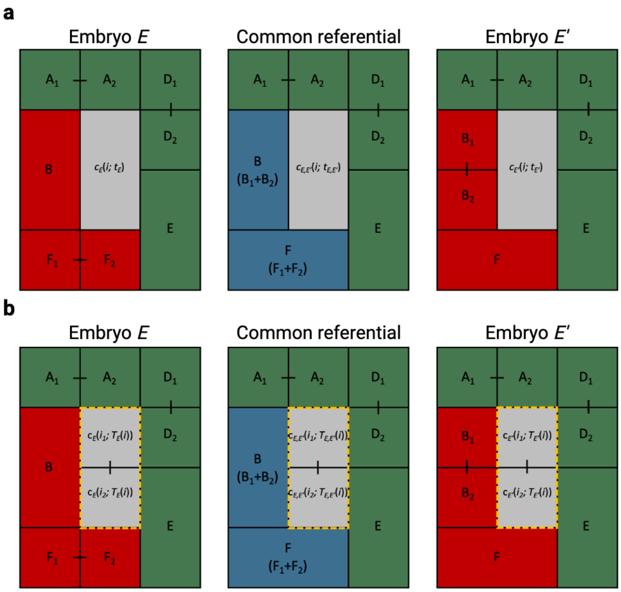


**Fig. S-12**: **Taking into account division heterochronies to compute local neighborhoods.** a) Illustration showing the neighborhood of a cell $c(i, t)$ (grey) in two different embryos, $E$ and $E'$ (see Supp. Methods B.7.1). To compute the neighborhood distance, sister cells present in only one of the two embryos (red) are virtually merged to define a common referential (blue). Sister cells present in the two embryos are in green. Black lines link sister cells. b) Illustration showing the neighborhood of the two daughter cells of a cell $c(i)$: $(c(i_1;\ T(i))$ and $c(i_2;\ T(i))$ (surrounded by yellow dotted lines) in two different embryos, $E$ and $E'$ (see Supp. Methods B.7.2). As with neighborhood distance, the division distance is computed in a common neighborhood referential (middle) using the closest common ancestors (blue).

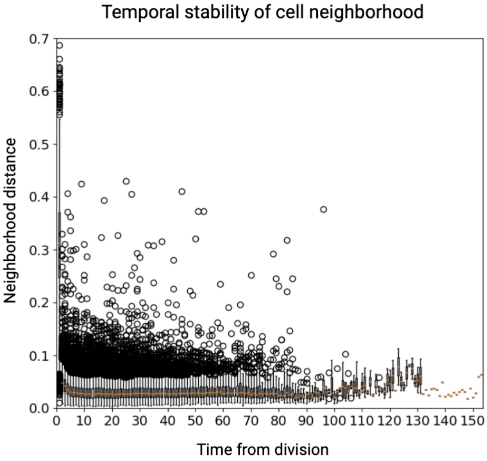


**Fig. S-13**: **Temporal stability of cell neighborhoods.** For each cell, the stability of its neighborhood was determined by calculating the neighborhood distance between two successive time points throughout the entire lifetime of the cell. Top, middle, and bottom lines in the box plot indicate $75^{th}$ percentile, median, and $25^{th}$ percentile of data, respectively.

| | | |
|---|---|---|
| Epidermis | | Head Epidermis |
| | | Lateral Tail Epidermis |
| | | Medio-Lateral Tail Epidermis |
| | | Midline Tail Epidermis |
| NS | | Anterior Ventral Neural Plate |
| | | Anterior-Dorsal Neural Plate |
| | | Posterior Ventral Neural Plate |
| | | Posterior-Lateral Neural Plate |
| | | Posterior-Dorsal Neural Plate |
| | | Germ Line |
| Mesoderm | | 1st Lineage, Notochord |
| | | 2nd Lineage, Notochord |
| | | Trunk Lateral Cell |
| | | Mesenchyme |
| | | Trunk Ventral Cell |
| | | 1st Lineage, Tail Muscle |
| | | 2nd Lineage, Tail Muscle |
| Endoderm | | Anterior Head Endoderm |
| | | Posterior Head Endoderm |
| | | 1st Endodermal Lineage |
| | | 2nd Endodermal Lineage |

**Fig. S-14: Larval tissue fates color table**

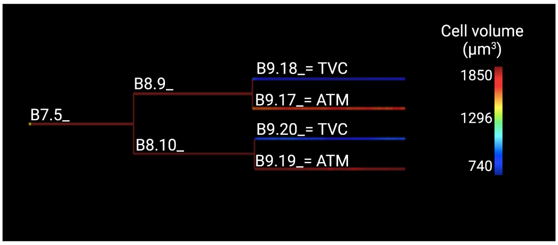


**Fig. S-15: Conklin name assignment to the cardiopharyngeal lineages.** Schematic lineage tree of the Phmamm-1-v2 B7.5_ cell with color code on branches indicating the cell volumes (scale bar on the right). The unequal divisions of B8.9_ in B9.18_ (smaller) and B9.17_ (bigger) and of B8.10_ in B9.20_ (smaller) and B9.19_ (bigger) allowed us to assign a name to the B8.9 and B8.10 daughter cells which were previously only named based on differential fate specification of the cardiopharyngeal progenitors (named Trunk Ventral Cells (TVC) or Anterior Tail Muscles (ATM) cells (**?**)).

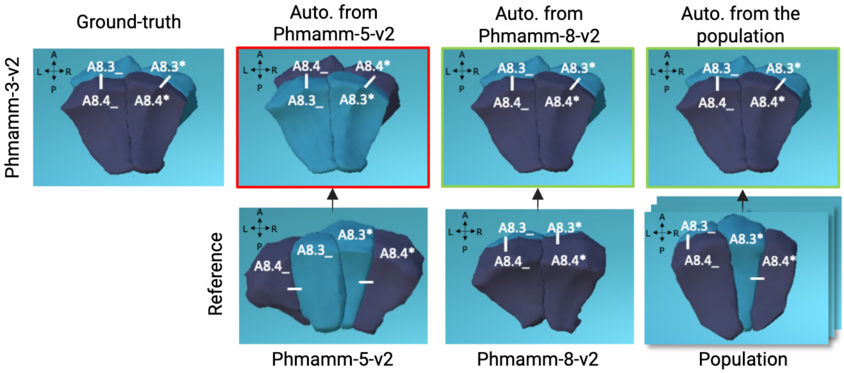


**Fig. S-16**: **Effect of the choice of the reference on the automated naming output.** A7.2 daughters of Phmamm-3-v2 (top left, manual corrected naming) are named differently when Phmamm-5-v2 or Phmamm-8-v2 are used as reference. The result of the automated naming from the Phmamm-v2 population of reference embryos (except Phmamm-3-v2) is also shown. Cell names are displayed and cells with the same name are color coded. Green and red boxes illustrate either a correct (green) or a wrong (red) automated naming of the A7.2 daughters in Phmamm-3 from the indicated reference. Vegetal views, Antero-Posterior (AP) and Left-Right (LR) axes are represented.

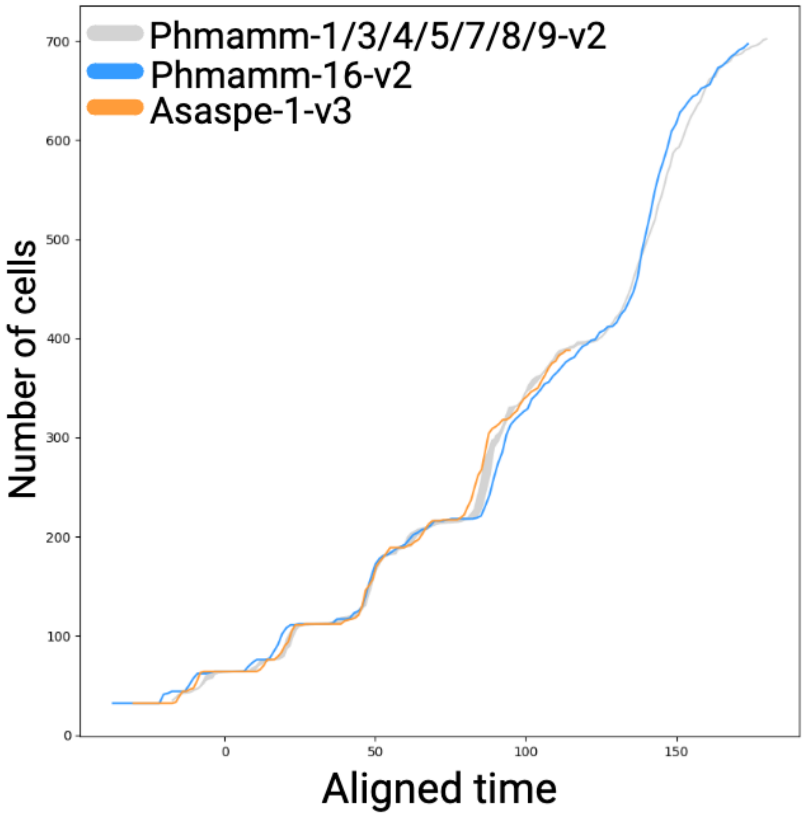


**Fig. S-17**: **Temporal alignment of reference embryos (Phmamm-1/3/4/5/7/8/9-v2) and the new ascidian segmented embryos: Phmamm-16-v2 (*Phallusia mammillata*) and Asaspe-1-v3 (*Ascidiella aspersa*).** The gray curve represents the space covered by all reference embryos (Phmamm-1/3/4/5/7/8/9-v2) in their cell number evolution after temporal alignment based on Phmamm1-v2. The blue and orange curves represent the evolution of cell number after temporal alignment on Phmamm1-v2 for Phmamm-16-v2 (blue) and Asaspe-1-v3 (orange).

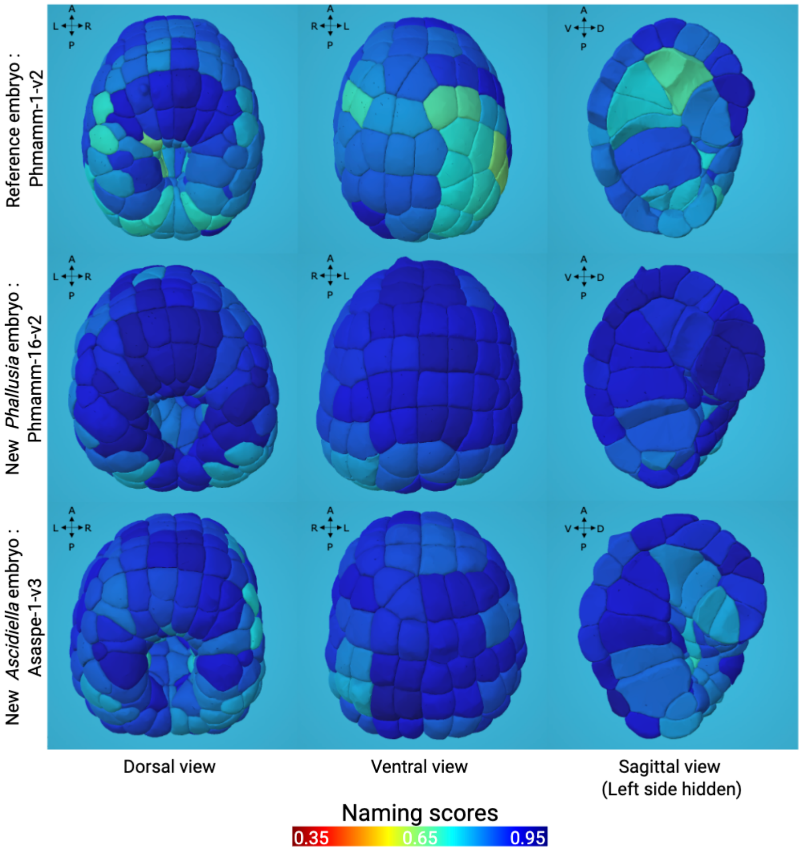


**Fig. S-18**: **Naming scores in the new segmented *Phallusia* and *Ascidiella* embryos compared to the reference embryo Phmamm-1-v2.** Dorsal (left), ventral (middle) and sagittal (right) views of a reference *Phallusia* embryo (Phmamm-1-v2, top), Phmamm-16-v2 (middle) and Asaspe-1-v3 (bottom) at the time step corresponding to 218-cells in Phmamm-1-v2 (see Figure S-17). Cells are color-coded according to the Naming score. Visualized with MorphoNet.

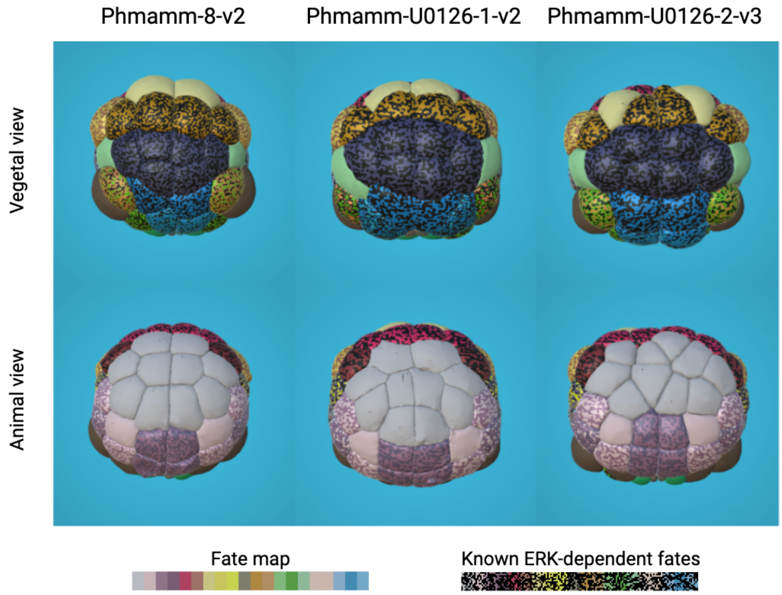


**Fig. S-19**: **Initial naming at stage 8 (64-cell) for the Phmamm-8-v2 reference embryo and the two MEK inhibitor-treated embryos.** Vegetative and animal views of a reference embryo (Phmamm-8-v2) and the two U0126-treated embryos, Phmamm-U0126-1-v2 (2µM U0126) and Phmamm-U0126-2-v3 (6µM U0126), at stage 8 (64-cell). Cells are color-coded according to the wild-type cell fates inherited from the assigned names (see Figure S-14 for the detailed tissue fates color table). In addition, multi-colored cells with black correspond to cells known to be induced by ERK signaling pathway activity or to have an ancestor that was induced by the ERK pathway earlier in development.

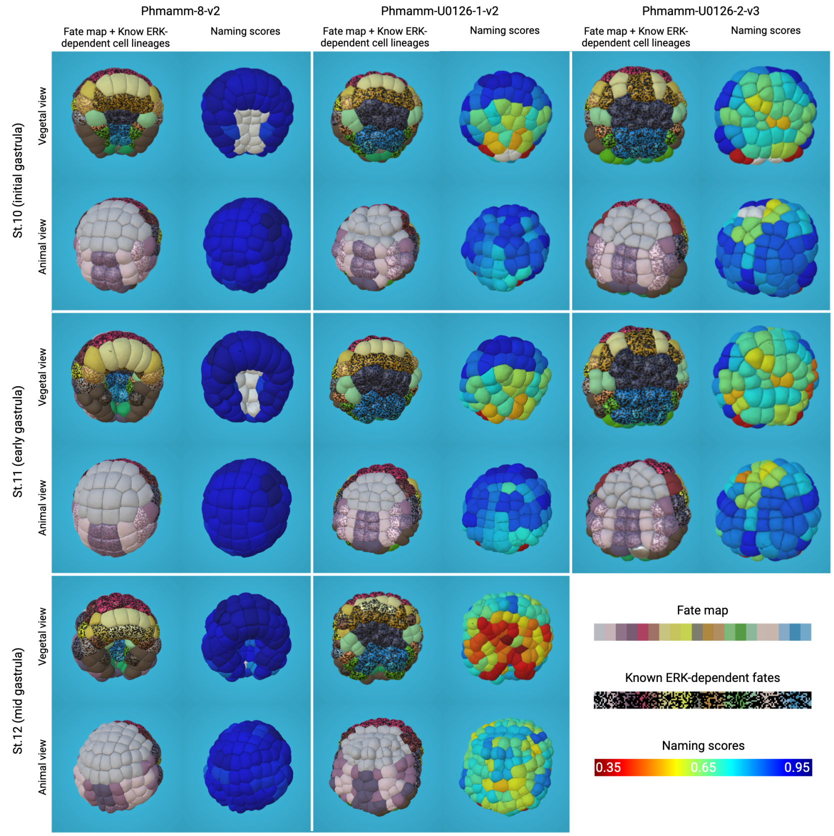


**Fig. S-20**: **Phenotypic analysis of the effects of MEK inhibition in *Phallusia* embryos using naming score.** Vegetative and animal views of a reference embryo (Phmamm-8-v2) and the two U0126-treated embryos, Phmamm-U0126-1-v2 (2µM U0126) and Phmamm-U0126-2-v3 (6µM U0126), at stages 10 (initial gastrula), 11 (early gastrula) and 12 (mid gastrula). For each stage, on the left: cells are color-coded according to the wild-type cell fates inherited from the assigned names (see Figure S-14 for the detailed tissue fates color table). In addition, multi-colored cells with black correspond to cells known to be induced by ERK signaling pathway activity or to have an ancestor that was induced by the ERK pathway earlier in development. On the right: cells are color-coded according to the naming score. Uncolored cells correspond to $7^{th}$ generation cells which have not yet divided, and whose name comes from initialization, and which therefore have no naming score calculated.
